# Molecular mechanism of TRPV2 channel modulation by cannabidiol

**DOI:** 10.1101/521880

**Authors:** Ruth A. Pumroy, Amrita Samanta, Yuhang Liu, Taylor E.T. Hughes, Siyuan Zhao, Yevgen Yudin, Kevin W. Huynh, Z. Hong Zhou, Tibor Rohacs, Seungil Han, Vera Y. Moiseenkova-Bell

## Abstract

Transient receptor potential vanilloid 2 (TRPV2) plays a critical role in neuronal development, cardiac function, immunity, and cancer. Cannabidiol (CBD), the non-psychotropic therapeutically active ingredient of *Cannabis sativa*, is a potent activator of TRPV2 and also modulates other transient receptor potential (TRP) channels. Here, we determined structures of the full-length TRPV2 channel in a CBD-bound state in detergent and in PI(4,5)P_2_ enriched nanodiscs by cryo-electron microscopy. CBD interacts with TRPV2 through a hydrophobic pocket located between S5 and S6 helices of adjacent subunits, which differs from known ligand and lipid binding sites in other TRP channels. Comparison between apo- and two CBD-bound TRPV2 structures reveals that the S4-S5 linker plays a critical role in channel gating upon CBD binding. The TRPV2 “vanilloid” pocket, which is critical for ligand-dependent gating in other TRPV channels, stays unoccupied by annular lipids, PI(4,5)P_2_, or CBD. Together these results provide a foundation to further understand TRPV channel gating properties and their divergent physiological functions and to accelerate structure-based drug design.

## INTRODUCTION

Transient receptor potential (TRP) channels play significant roles in human physiology and facilitate permeation of essential ions (Na^+^, Ca^2+^) through the plasma membrane^1,2^. Transient receptor potential vanilloid 2 (TRPV2) belongs to the thermoTRPV subfamily of TRP channels (TRPV1-TRPV4), yet TRPV2 is insensitive to both vanilloids and heat^3^. It has been the least studied among TRPVs due to the lack of specific pharmacological agonists or antagonists^4^. Cannabidiol (CBD), a natural product of the *Cannabis sativa* plant, is a potent TRPV2 agonist^4^ which has been recently used to demonstrate the important role of TRPV2 in the inhibition of glioblastoma multiforme cell proliferation^5–9^. These findings place TRPV2 on the list of important anti-tumor drug targets^5–9^.

Cannabinoids and cannabinoid analogs have been reported to activate a variety of TRP channels, including TRPV1-4^4,10,11^ and TRPA1^12^. CBD activates TRPV2 most potently, with an EC_50_ of 3.7 μM for rat TRPV2^4^, making it an excellent candidate for the investigation of TRP channel modulation by CBD using cryo-electron microscopy (cryo-EM). Additionally, the modest activation of other TRP channels by CBD suggests that a cannabinoid binding site could be conserved within this family of channels^4,10–12^. Understanding the molecular mechanism of CBD activation of functionally diverse TRP channels could allow us to gain insight into the gating mechanisms of these channels and develop novel modulators.

The TRPV channel subfamily members (TRPV1-TRPV6) have strong structural homology^13^, yet the effects of lipids on these channels are widely divergent^14–18^. Structural studies of members of this family have shown that both modulating and structural lipids can bind the “vanilloid” pocket located in the transmembrane domain (TMD) between the S3 and S4 helices and the helical S4-S5 linker of each monomer^15–17^. This pocket has also been shown to be a drug binding site in some TRPV channels and different occupants of this pocket can transmit conformational changes into the pore which affect gating^15,16^. This “vanilloid” pocket does not have very strong lipid density in currently available structures of TRPV2^18–20^, TRPV3^21,22^ and TRPV4^23^ channels, while annular or modulating lipids have strong density in this pocket in TRPV1^15^, TRPV5^16^ and TRPV6^17^ channels. One lipid of interest for this study is phosphatidylinositol 4,5-bisphosphate (PI(4,5)P_2_), which is a known modulator of all TRPV channels^14^, but it is uncertain whether the mechanisms of modulation are conserved among these closely related channels^14^. PI(4,5)P_2_ has been shown to both activate^24^ and inhibit TRPV2^18^, but the exact molecular mechanism is poorly understood^14^. On the other hand, the PI(4,5)P_2_ activation mechanism of TRPV5 and TRPV6 is conserved^25^ and a novel PI(4,5)P_2_ binding site has recently been revealed using cryo-EM^16^.

To investigate the effect of CBD and lipid binding on TRPV2 channel gating, we determined the cryo-EM structure of the full-length TRPV2 channel in a CBD-bound state at 4.3 Å resolution in detergent and at 3.5 Å resolution in PI(4,5)P_2_ enriched nanodiscs. The CBD-bound TRPV2 structure in nanodiscs clearly revealed that CBD interacts with the channel through a novel hydrophobic ligand binding pocket located between the S5 and S6 helices. CBD binding to the channel in nanodiscs induced an α- to π-helix transition in the S4-S5 linker, however this conformational change did not produce significant changes in the pore, suggesting that we may have captured TRPV2 in either an intermediate agonist-bound state or a desensitized state. The TRPV2 “vanilloid” binding pocket remained unoccupied by either CBD or PI(4,5)P_2_, despite enrichment of the lipid bilayer with soluble PI(4,5)P_2_. These results imply that TRPV2 physiological activity may not depend on tightly bound annular or modulating lipids in this pocket, which are essential for the function of some other TRPV channels. Despite lacking clear density for CBD, the CBD-bound TRPV2 structure in detergent revealed that CBD can induce conformational changes in the S4-S5 linker of the channel that allowed us to capture TRPV2 in a partially open state. Together these results provide new information on TRPV2 ligand-dependent gating and this knowledge could be further explored for the design of novel TRP channel therapeutics.

## RESULTS

### Detergent-solubilized TRPV2 in CBD-bound state

To determine the structure of TRPV2 in a CBD-activated state, we first used detergent solubilized full-length rat TRPV2 protein^19^ and incubated it for 30 min with 30μM CBD before freezing it in vitreous ice. This corresponds to ten times the previously reported EC_50_^4^ and the reported incubation time with various ligands for truncated TRPV1^15,26^. The cryo-EM images revealed a monodisperse distribution of detergent-solubilized TRPV2 protein in vitreous ice in the presence of 30μM CBD (Supplementary Figure 1). After 2D classification, the best particles were subjected to 3D auto-refine and 3D classification in RELION^27^, which yielded one distinct 3D class with four-fold symmetry. This class was further classified and refined to yield a final TRPV2 channel in a CBD-bound state at 4.3 Å resolution (Supplementary Figure 1, Supplementary Figure 2, Supplementary Table 1). The higher-resolution features of this class in the transmembrane region of the channel (~4.0 Å) allowed us to build a model for this map and place sidechains in the transmembrane region of the channel (Supplementary Figure 1, Supplementary Figure 3). As in previous cryo-EM structures of TRPV2^18–20^, it forms a homo-tetramer featuring six transmembrane helices (S1-S6) spanning the transmembrane domain (TMD) with six ankyrin repeat domains (ARDs) splayed out like a pinwheel on the cytoplasmic face of the protein, with the ARDs of adjacent monomers glued together through a β-sheet region (Supplementary Figure 2). S1-S4 form a bundle, while S5, S6 and the pore helix extend outwards to domain swap with adjacent monomers and form the pore. Despite working with the full-length rat TRPV2 protein, our map lacks density for the ~30 residues that make up the pore turret between the top of S5 and the pore helix, suggesting that the pore turrets form a flexible loop.

To have a better comparison than the currently available apo states of TRPV2 ^18–20^, we processed the original movies for apo full-length rat TRPV2 in detergent from scratch using RELION 3.0^28^, which we had collected for our previously published apo full-length rat TRPV2 structure^19^. New algorithms for motion correction^29^, CTF correction^30^ and autopicking^28^ yielded an improved selection of particles from the original movies. This new dataset, which cannot be compared to the previously published dataset^19^, yielded a map at 4.2 Å resolution (Supplementary Figure 4, Supplementary Figure 5). We were able to build a model for this map and place sidechains in most of the transmembrane region of the channel, which had local resolution at ~4.0 Å (Supplementary Figure 4, Supplementary Figure 6).

Comparison between TRPV2 in the apo state at 4.2 Å resolution and the CBD-bound state at 4.3 Å did not show obvious density for a drug but did reveal that the channel adopted a partially open conformation (Figure 1A-C). It is occluded at the lower gate (Met645) (Figure 1C), which would prevent ion permeation through the channel (Figure 1C). On the other hand, the application of 30μM CBD induced global rearrangements, subtly pulling the tops of S5, S6 and the pore helix outwards (Figure 1D).

**Figure 1.**
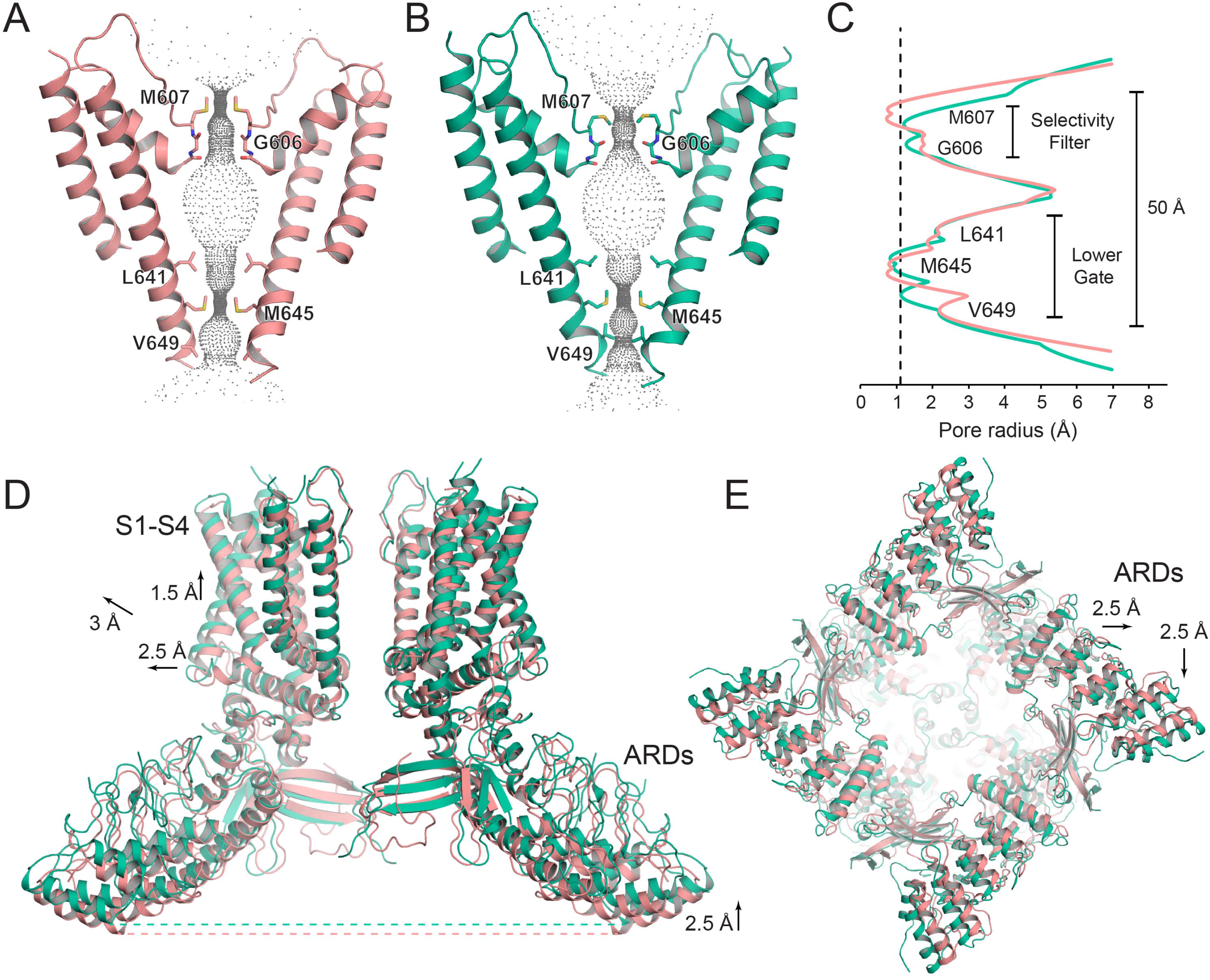
Global changes associated with CBD binding. (**A-B**) Cartoon dimer pore representation of (A) apo TRPV2 and (B) CBD-bound TRPV2 in detergent. Grey dots indicate the diameter of the ion conduction pore. Constriction residues are labeled and shown as sticks. (**C**) Graphical representation of the radius of the pore as a function of the distance along the ion conduction pathway. The radius for the pore of apo TRPV2 is shown in salmon and the CBD-bound TRPV2 in detergent pore is shown in green. The dotted line indicates the radius of a dehydrated calcium ion. (**D**) Overlay of opposing dimers of apo (salmon) and CBD-bound TRPV2 in detergent (green). Arrows indicate direction of movement and are labeled with the distance moved. (**E**) Bottom view of the overlaid tetramers of apo (salmon) and CBD-bound TRPV2 in detergent (green). Arrows indicate direction of movement and are labeled with the distance moved.

Beyond the changes at the pore, the addition of 30μM CBD also induced dramatic conformational changes to S1-S4, the TRP helix, and the ARDs, with an overall RMSD between apo and CBD-bound TRPV2 in detergent of 2.5 Å (Figure 1D, E). The S1-S4 bundle and TRP helix move together outwards by ~2.5 Å and upwards by ~1.5 Å, for a total translation of ~3 Å (Figure 1D). The larger outward shift of the S1-S4 bundle creates space for the upper parts of S5, S6 and the pore helix to move slightly outwards, which leads to the opening of the selectivity filter (Figure 1A, B). The outward shift of the TRP helix also shifts the C-terminal end of the ARD region outward while the N-terminal end of the ARDs is anchored by an interaction with the β-sheet region of an adjacent monomer, causing the ARDs to pivot. This moves the most C-terminal portion of the ARDs outwards by ~2.5 Å in the same direction as the TRP helix, but with less than 0.5 Å upward movement, while the N-terminal portion of the ARDs move upwards by ~2.5 Å and outwards by ~2.5 Å perpendicular to the movement of the TRP helix (Figure 1D, E). In the apo TRPV2 structure, the most C-terminal residues visible (Glu716-Pro729) wrap around the bottom of the β-sheet region, making contacts with ARDs 2-4 of the adjacent monomer. Interestingly, in the CBD-bound structure, the most C-terminal residues move away from the tail and are only visible up to Leu719, but there is still density for ~16 residues in a V-shaped loop between the β-sheet region and ARDs 1-3 of the adjacent monomer (Figure 2A). This density has no continuity with the rest of the map, but due to the distance from the C-terminus we predict that this density should be attributed to unknown residues from the N-terminus.

**Figure 2.**
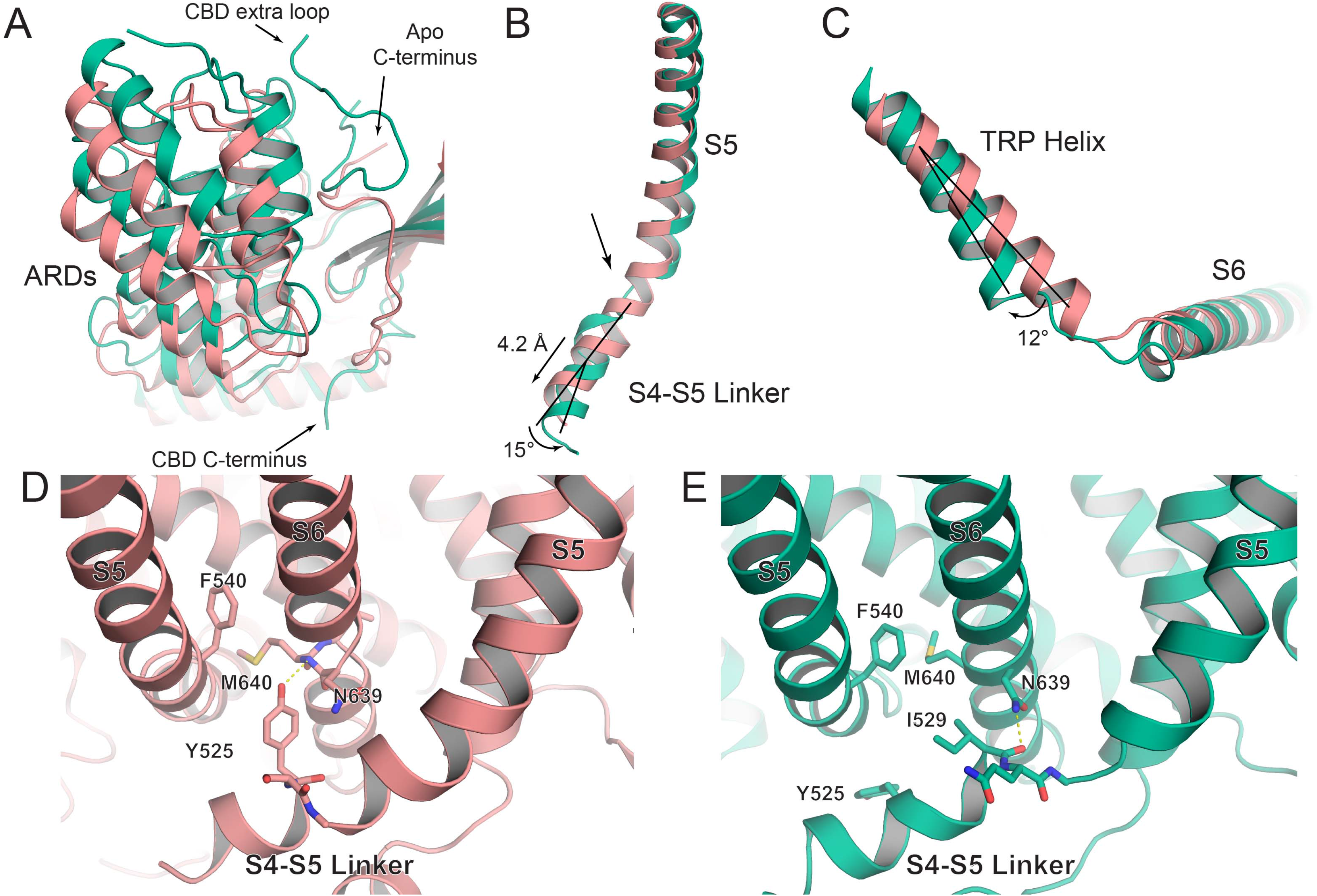
Conformational changes associated with CBD binding. (**A**) An overlay of a single ARD from the apo (salmon) and CBD-bound TRPV2 in detergent (green) models. Arrows indicate loops that differ between the two models. (**B**) The S4-S5 linker, S5 helix and S6 helix of apo (salmon) and CBD-bound TRPV2 in detergent (green). Black lines indicate the trajectory of the S4-S5 linker. The measurements for the lateral movement and angle pivoted are labeled. The area of helix breakage is indicated by a black arrow. (**C**) The TRP helix and lower S6 helix of apo (salmon) and CBD-bound TRPV2 in detergent (green). Black lines indicate the trajectory of the TRP helix. The measurement for the angle pivoted is labeled. (**D-E**) Area of helix breakage for (D) apo TRPV2 and (E) CBD-bound TRPV2 in detergent. Residues of interest are labeled and shown as sticks.

The origin of these large movements seems to be the S4-S5 linker and S5 helix. The bottom of the S5 helix of the CBD-bound structure bows outwards from the pore and the S4-S5 linker shifts away from S5 by 4.2 Å and rotates outwards by 15°, causing a break in the helix at residues Gln530, Lys531, and Val532 (Figure 2B). In addition to pushing the S1-S4 and TRP helix outwards, this movement also rotates the N-terminal end of the TRP helix by 12° (Figure 2C). The combined outward movement and rotation of the TRP helix unravels the first 3 residues of helix adjacent to S6, which also allows for a slight change in the rotation of the bottom of the S6 helix, bringing Val649 further into the pore (Figure 1B). This movement seems to be initiated by the large outward movement of S5. In apo TRPV2 in detergent, Tyr525 on the S4-S5 linker fits into a pocket between S5 and S6 of an adjacent monomer (Figure 2D). In CBD-bound TRPV2 in detergent, the outward bowing of S5 moves Phe540 2 Å into this pocket, pushing Tyr525 out of the pocket so that it is partially exposed to solvent on the cytoplasmic face of the protein (Figure 2E). Additionally, this puts Asn639 in a position to form a hydrogen bond with the backbone carbonyl of residue Ile529. Based on all of these conformational changes, we suggest that this may represent a partially open state of the channel.

### CBD-bound TRPV2 in PI(4,5)P_2_ enriched nanodiscs

To improve our structure and identify the CBD binding site, we reconstituted full-length rat TRPV2 into nanodiscs in the presence of 400μM PI(4,5)P_2_ dioctanoyl (diC8). 100μM diC8 PI(4,5)P_2_ was reported to mostly restore the activity of desensitized TRPV2 in PI(4,5)P_2_-depleted membranes^24^, so we used four times that amount to ensure saturation, a concentration which had also been sufficient to visualize diC8 PI(4,5)P_2_ in our recently published TRPV5 structure^16^. We also incubated our TRPV2 sample with 100μM CBD, which corresponds to thirty times the previously reported EC_50_^4^ in order to saturate the protein with the ligand. We exposed TRPV2 to these modulators both during reconstitution into nanodiscs and during a final incubation before preparing grids. Cryo-EM images revealed a monodispersed distribution of TRPV2 protein in vitreous ice (Supplementary Figure 7). After 2D classification, the best particles were subjected to 3D auto-refine and 3D classification in RELION^28^, which again yielded one distinct 3D class with four-fold symmetry. This class was further classified and refined to yield a final map with clear density for CBD at 3.5 Å resolution (Supplementary Figure 7, Supplementary Figure 8). Due to the high quality of the map (Supplementary Figure 9), we were able to build an atomic model of TRPV2 and unambiguously position the CBD ligand and sidechains throughout the TMD, but found no density that could be attributed to PI(4,5)P_2_ and again did not observe any density for the TRPV2 pore turrets.

The comparison between the CBD-bound TRPV2 pore in nanodiscs and the apo TRPV2 pore in detergent revealed that despite some differences in sidechain positions, both pores are still closed at both the selectivity filter and lower gate (Figure 3A, B). We were able to identify the CBD binding pocket and observe modest conformational changes in the S4-S5 linker, S1-S4 bundle, TRP helix, and ARDs, with an overall RMSD of 1 Å. In comparison to the ~3 Å outward expansion of the S1-S4 bundle in the CBD-bound structure in detergent (Figure 1D), the same region of the CBD-bound structure in nanodiscs expands outwards in a similar direction, but only by ~1 Å. This suggests that we could have trapped TRPV2 in a CBD-bound intermediate state, or due to the timescale of CBD application before grid preparation and the CBD concentration above EC_50_, we could have trapped the channel in a desensitized state.

**Figure 3.**
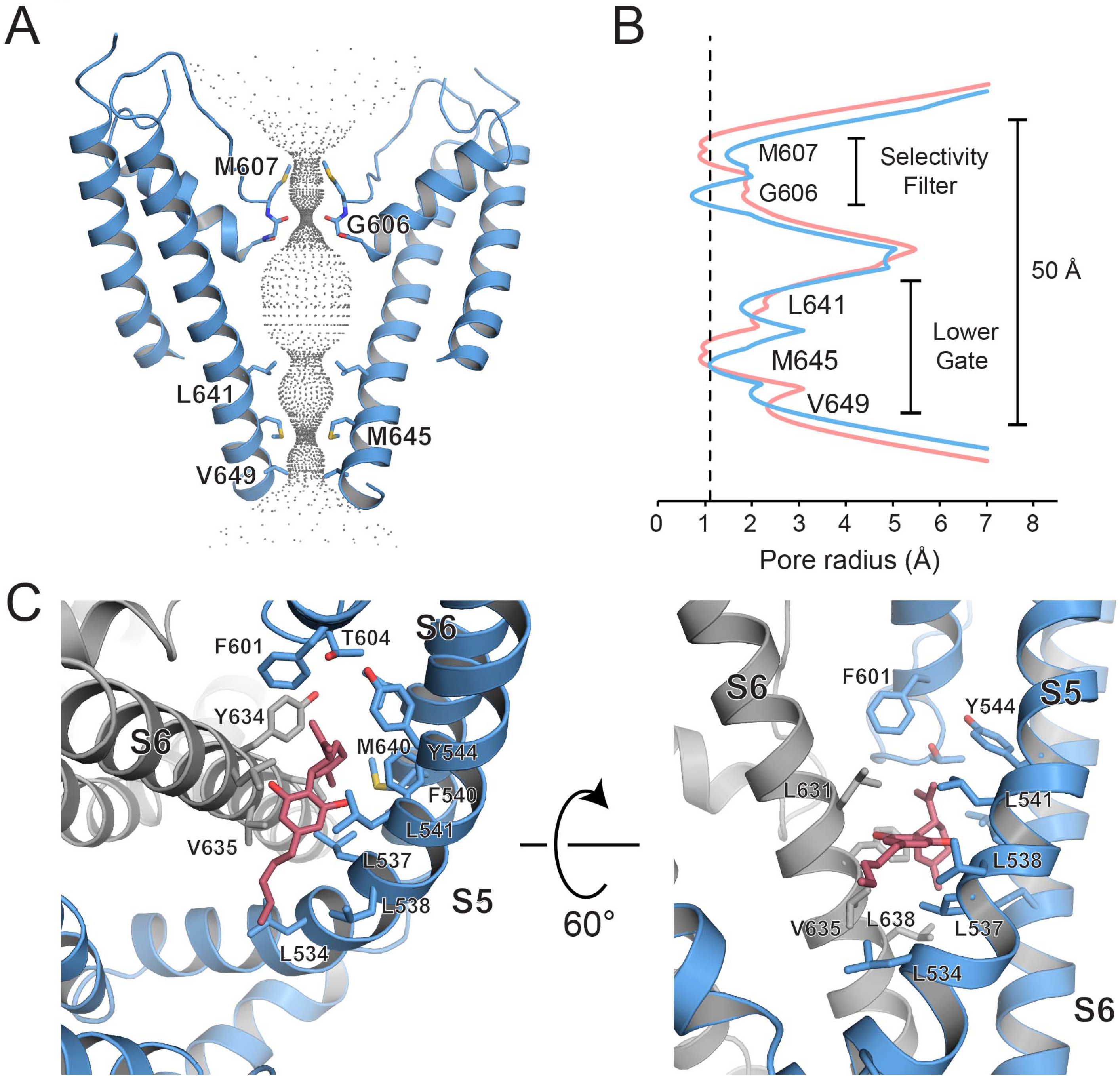
CBD-bound TRPV2 in PI(4,5)P_2_ enriched nanodiscs. (**A**) Cartoon dimer pore representation of CBD-bound TRPV2 in nanodiscs. Grey dots indicate the diameter of the ion conduction pore. Constriction residues are labeled and shown as sticks. (**B**) Graphical representation of the radius of the pore as a function of the distance along the ion conduction pathway. The radius for the pore of CBD-bound TRPV2 in nanodiscs is shown in blue and the apo pore is shown in salmon. The dotted line indicates the radius of a dehydrated calcium ion. (**C**) Model representation of the CBD binding pocket in the CBD-bound TRPV2 in nanodiscs structure. The S5 and S6 helices of TRPV2 are shown as blue and grey cartoons, respectively. CBD is shown as pink sticks. Residues of interest are labeled and represented as sticks.

The CBD binding site is located between the S5 and S6 helices (Supplementary Figure 10) of adjacent TRPV2 monomers and is lined with hydrophobic and aromatic residues, including Leu631, Tyr634, Val635 and Leu638 on the S6 helix of one monomer and Leu534, Leu537, Leu538, Phe540, Leu541, and Tyr544 on the S5 helix of an adjacent monomer (Figure 3C). The S6 of this adjacent monomer also contributes contacts at Leu637 and Met640. The pore helix of the adjacent monomer forms the cap of the pocket, blocking CBD from entering the pore with residues Phe601 and Thr604 (Figure 3C). CBD is composed of a hydrophobic head group, a middle aromatic ring with two hydroxyl groups, and a 5-carbon tail (Supplementary Figure 10). The hydrophobic head group enters furthest into the pocket, pushing Phe540, Tyr634 and Leu638 outwards from the CBD-binding pocket relative to their positions in the reprocessed apo TRPV2 model (Supplementary Figure 10). The hydroxyl groups of the middle region fit between turns of the alpha helices on either side, coordinating with the backbone hydrogen bonds between Leu631 and Val635 on the S6 of one monomer and Leu537 and Leu541 on the S5 of the adjacent monomer, and pushing Leu537 down and out of the way relative to its position in apo TRPV2 (Figure 3C). The 5-carbon tail has weaker density (Supplementary Figure 10, Supplementary Figure 11), indicating flexibility where the CBD tail meets lipids in the nanodisc bilayer.

The density for CBD in this binding pocket in the CBD-bound TRPV2 in nanodiscs structure is prominent and clearly present in both half maps (Supplementary Figure 11). As of the time of writing this manuscript, no other TRPV channel structures^15–23^ were reported to have lipid density in this pocket, so in combination with the good fit of the drug to the density, we are confident in assigning it to CBD. With the identification of the CBD binding pocket in the nanodisc structure, we re-examined the map of CBD-bound TRPV2 in detergent at the same pocket (Supplementary Figure 11). In the half maps for CBD-bound TRPV2 in detergent there is some very weak extra density in this pocket, while none appears at the same level in the apo TRPV2 in detergent maps (Supplementary Figure 11). More strikingly, the binding pocket in the CBD-bound TRPV2 in detergent map had expanded relative to the apo TRPV2 in detergent map to the size required to accommodate a molecule of CBD (Figure 4A). The distance between Tyr634 and Phe540 expanded by 1.5 Å between the apo and the CBD-bound TRPV2 in detergent models, from 5.7 Å to 7.2A, almost the same distance between these residues in the CBD-bound TRPV2 in nanodiscs structure (Figure 4A). It’s interesting to note that the two CBD-bound structures expand the binding pocket in different ways (Figure 4A, B, C). In the nanodisc structure, S5 bows out slightly, moving Phe540 outwards, but Tyr634 also turns away from the pocket (Figure 4B). By contrast, in the detergent structure Tyr634 does not move, but S5 moves outwards more dramatically, moving Phe540 along with it (Figure 4C). This suggests that CBD is in fact present, if not resolved, in the CBD-bound TRPV2 in detergent structure. The absence of clear density for CBD in the CBD-bound detergent structure is likely due to a combination of factors, including the lower resolution of this data and occupancy of the pocket (Supplementary Figure 11). The highly hydrophobic nature of CBD would have caused it to accumulate in the nanodiscs, effectively increasing its local concentration even further in the CBD-bound TRPV2 in nanodiscs sample. A recent paper from the Lee group^22^ reported a similar situation for their structures of TRPV3 in the presence of 2-APB: although they were able to observe structural changes to the channel upon exposure to the activator, they were not able to identify density for the 2-APB^22^.

**Figure 4.**
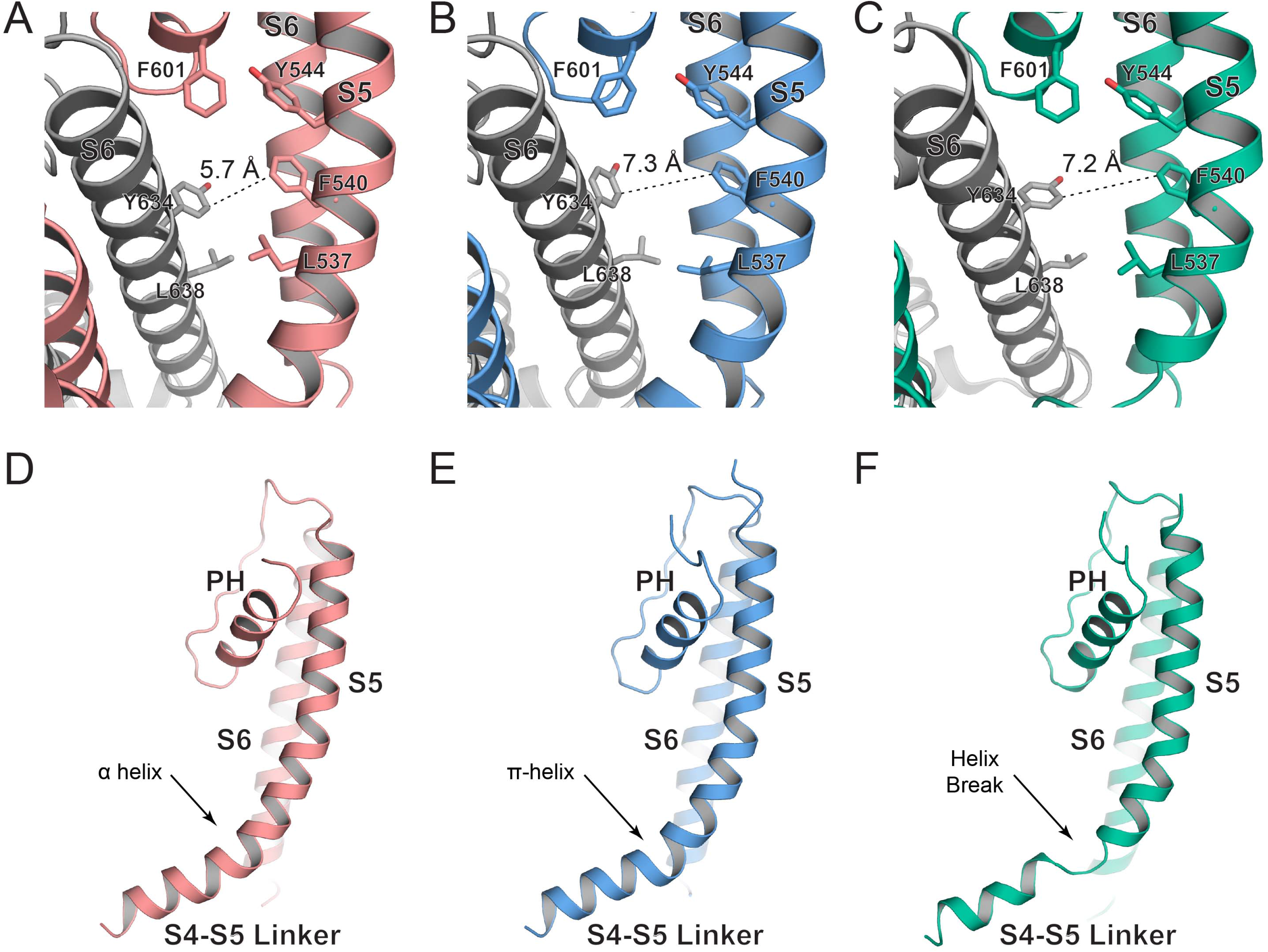
Conformational changes in the CBD binding pocket. (**A-C**) Model representations of the CBD binding pockets in the (A) apo TRPV2 (B) CBD-bound TRPV2 in nanodiscs (C) CBD-bound TRPV2 in detergent structures. Residues of interest are labeled and shown as sticks. The distance between Tyr634 and Phe540 for each model is indicated by the dotted black line and labeled. (**D-F**) S4-S6 helices of the (D) apo TRPV2 in detergent (E) CBD-bound TRPV2 in nanodiscs (F) CBD-bound TRPV2 in detergent structures. The arrow indicates the region in the S4-S5 linker that undergoes conformational rearrangement.

The changes observed at the CBD binding pocket in both CBD-bound structures are transmitted to conformational changes at the S4-S5 linker, which acts as a hinge to move the S1-S4 bundle, TRP helix, and ARDs. In the apo TRPV2 in detergent structure, the S4-S5 linker and S5 form a continuous, smoothly curved α- helix (Figure 4D). The slight outward movement of S5 in CBD-bound TRPV2 in nanodiscs induced a transition to a π-helix at residues Lys531-Ile533, forming a sharp bend between the S4-S5 linker and S5 (Figure 4E). The larger outward bulge of S5 in CBD-bound TRPV2 in detergent caused a total break in the helix at residues Lys531-Ile533, allowing the remaining helix of the S4-S5 linker to rotate and move away from S5 by ~5.5 Å (Figure 4F). Although the movement of S5 in the CBD-bound TRPV2 in nanodiscs structure pushed Phe540 into the pocket between S5 and S6 above the S4-S5 linker, this did not expel Tyr525 from the pocket as in the CBD-bound structure in detergent (Supplementary Figure 12). Instead, Tyr525 forms a hydrogen bond with the backbone amide of Met640, as recently seen in the apo TRPV3 structure from the Lee group between the homologous residues Tyr575 and Met672 (PBD 6MHO) (Supplementary Figure 12). As these bonds are broken in their 2-APB sensitized TRPV3 structure featuring a π-helix in S6 (PDB 6MHS), the Lee group^22^ suggests that this interaction stabilizes the α-helix of S6 preventing it from forming a π-helix, which is implicated in channel opening^17,21,22^ (Supplementary Figure 12).

In order to verify that this density is CBD and that these identified residues compose the CBD-binding pocket in TRPV2, we mutated key amino acids and performed Ca^2+^ imaging experiments. Due to the location of this pocket within the TMD, mutations to polar or charged residues were not feasible. We predicted that mutating Leu541, Leu631 and Val635, residues at the edge of the binding pocket, to bulky hydrophobic residues like phenylalanine may affect the entrance to the CBD binding pocket (Figure 3, Supplemental Figure 10). We found that in HEK293 cells transfected with the double mutant Leu541Phe-Leu631Phe, application of 20 and 50 μM CBD induced larger Ca^2+^ responses than in cells transfected with the wild type TRPV2 (Supplementary Figure 13). The Val635Phe mutant had severely impaired function, it only showed responses to CBD in a small percentage of cells (<5%), which were similar to those in control cells not transfected with TRPV2 (data not shown). Together, our results provide compelling evidence that this identified pocket is involved in CBD binding and channel modulation.

### TRPV2 lipid interactions

Many previous TRPV cryo-EM maps have shown strong lipid densities in hydrophobic pockets in the TMD^15–17,21,22^. The “vanilloid” pocket of TRPV1 is occupied by phosphatidylinositol (PI) in the apo structure in nanodiscs^15^ (Figure 5A), and TRPV5^16^ and TRPV6^17^ both report obvious densities for annular lipids in this pocket in proteins prepared in both detergents and nanodiscs. Both sets of recent TRPV3 structures have prominent lipid densities sitting above the “vanilloid” pocket between the tops of three helices: S5 and S6 of one monomer and S4 of an adjacent monomer^21,22^. Most TRPV cryo-EM maps also have pronounced density for some lipid nestled in the pocket formed by the S1, S2 and TRP helices^15–17,19–22^. The persistence of these lipids through purification processes suggests high affinity and potentially even a structural role, an idea supported by the alteration of pore structure after mutations to lipid binding pockets recently seen for TRPV3^21^ and TRPV6^17^.

**Figure 5.**
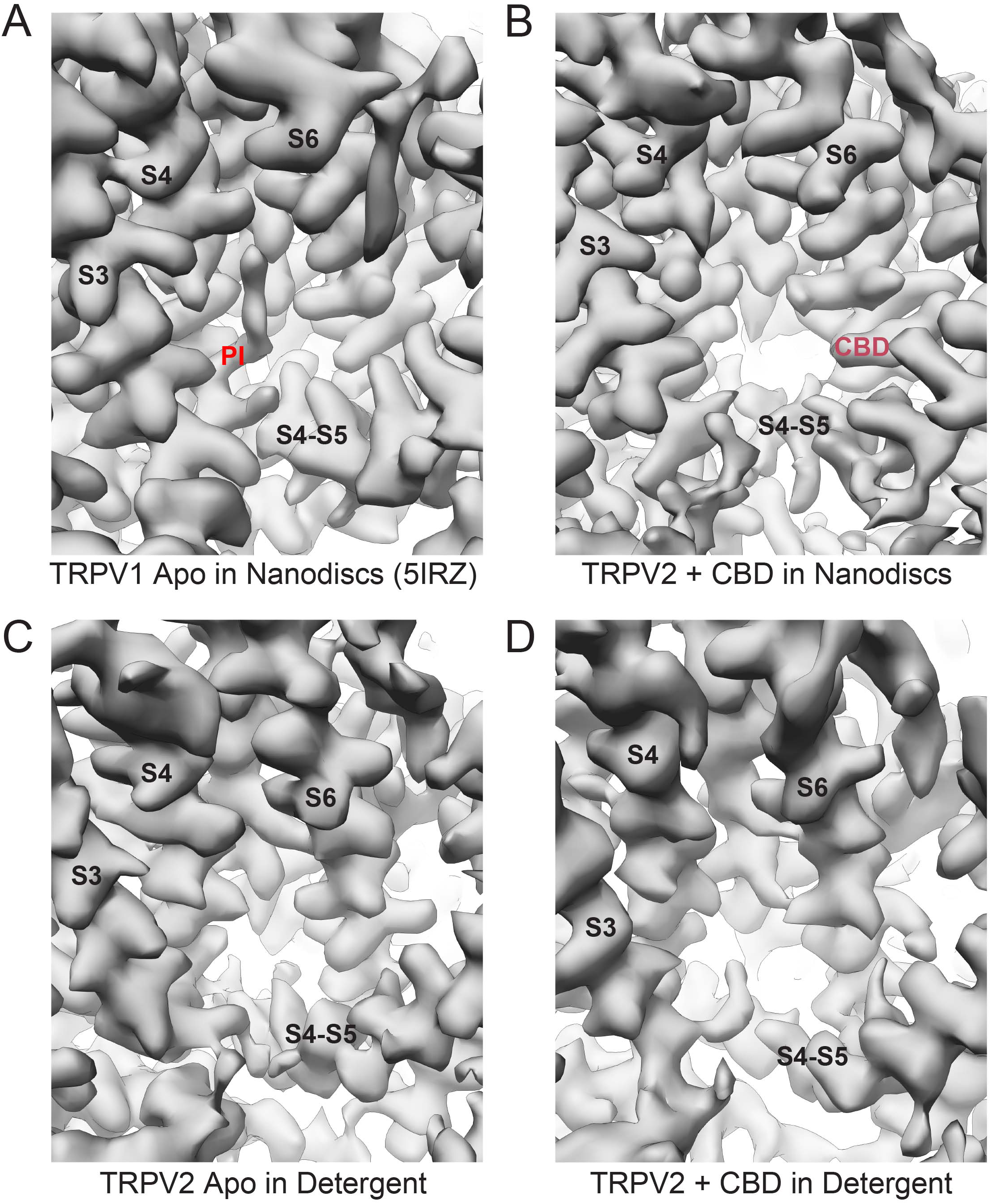
Vanilloid binding pocket. The vanilloid pocket of (**A**) apo TRPV1 in nanodiscs (PDB 5IRZ), (**B**) CBD-bound TRPV2 in nanodiscs, (**C**) apo TRPV2 in detergent and (**D**) CBD-bound TRPV2 in detergent. The models for the helices that constitute the pockets are shown as ribbons and overlaid with their respective cryo-EM densities shown as grey surfaces. Arrows indicate the location of densities in these pockets.

In contrast to these other TRPV cryo-EM maps^15–17,21,22^, previous cryo-EM maps of TRPV2 have consistently lacked obvious lipid density in the vanilloid pocket^18–20^. The improved full-length TRPV2 maps in detergent and nanodiscs presented here continue this trend (Figure 5B, C, D) and the addition of PI(4,5)P_2_ to TRPV2 during nanodisc reconstitution did not reveal PI(4,5)P2 in the “vanilloid” binding site (Figure 5B). Nor could we identify PI(4,5)P_2_ density in the binding pocket observed in TRPV5 located between the N-linker, S4-S5 linker, and S6 helix that has recently been revealed by cryo-EM^16^. In contrast, all three of these maps have pronounced density in the pocket formed by S1, S2 and the TRP helix, density which is the same shape and size as the lipid density in this pocket in the apo TRPV1 in nanodiscs structure^15^, identified as phosphatidylcholine (PC) (Supplementary Figure 14). We also observed two additional densities for lipids around the periphery of the TMD of the CBD-bound TRPV2 structure in nanodiscs (Supplementary Figure 8). Together, this data suggests that while full-length rat TRPV2 does have strong affinity for some lipids, it does not have strong interaction with PI(4,5)P_2_.

## DISCUSSION

Here, we have presented an improved full-length apo TRPV2 structure and CBD-bound TRPV2 structures in detergent and in PI(4,5)P_2_ enriched nanodiscs. We were able to identify the novel CBD binding site in the TRPV2 channel and determine conformational changes in TRPV2 upon CBD binding. TRPV2 interacts with CBD through a hydrophobic pocket located between S5 and S6 helices of adjacent subunits, which is conserved among TRPV channels (Figure 6). CBD binding induces the outward movement of S5, which leads to the expulsion of Tyr525 from the binding pocket between S5 and S6. This combined with the transition at the S4-S5 linker from α- to π-helix and eventually to a total break in the S4-S5 linker helix at residues Lys531-Ile533 causes larger conformational changes in the ARDs, S1-S4 bundle and TRP helix and pore of the channel that may allow for TRPV2 to open (Figure 6). Although none of the structures reported here appear to be in the fully open conformation, comparison of our current structures to the recently published open state of rat TRPV2 (PDB 6BO4) is not possible due to the quality of this open state map^18^. In spite of the resolution reported for the open state structure (PDB6BO4)^18^, the cryo-EM density is of poor quality and appears to be at a lower resolution than reported, which does not allow for precise identification of helical register or side chain placement (Supplementary Figure 15). As such comparing this structure to the ones presented in this manuscript would not be appropriate.

**Figure 6.**
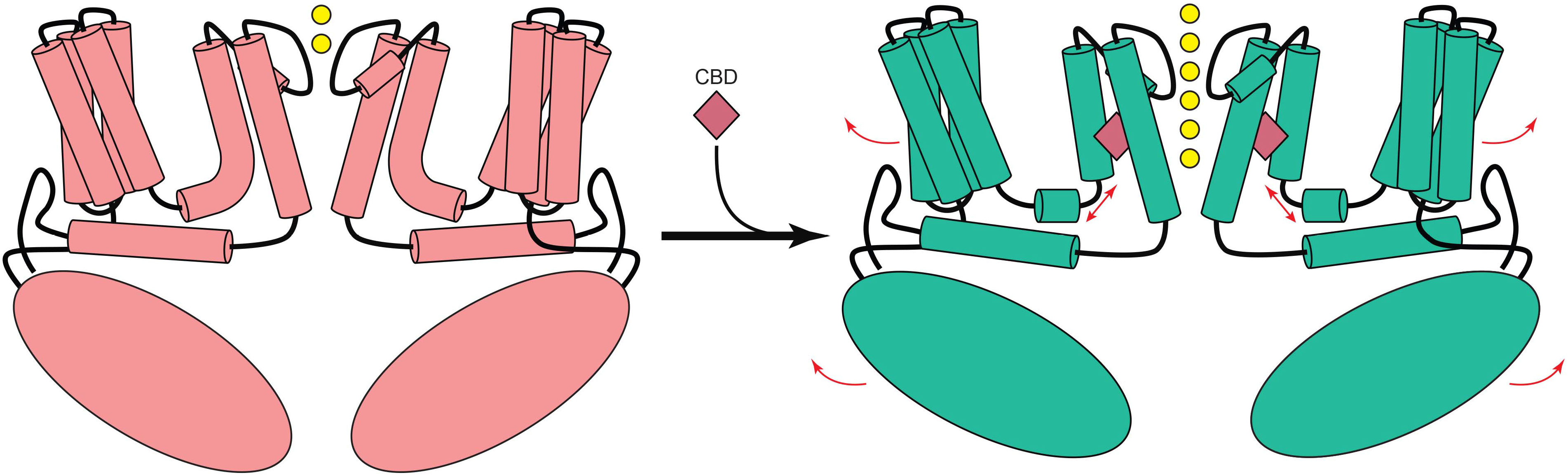
TRPV2 modulation by CBD. Schematic summary of the conformational changes observed due to CBD binding. The pink diamond represents molecular CBD. Arrows indicated movement of the channel. Yellow circles represent ions.

TRPV channels share ~40% sequence homology^13^ and thermoTRPV channels (TRPV1-TRPV4) have all been shown to have varying levels of modulation by CBD^4,10,11^. CBD has the largest positive effect on TRPV2^4^, and the structures of TRPV2 presented in this paper, along with those of other TRPV channels may give some insight into that preference despite very high sequence homology among the residues of the CBD binding pocket (Supplementary Figure 16). Several structures of thermoTRPV channels TRPV1^15^ and TRPV3^21,22^ show a π-helix near the top of S6, at the level of the CBD binding pocket. The presence of this π-helix has been suggested to be dynamic and potentially involved in pore opening, and the recent TRPV3 structures from both the Lee and Sobolevsky groups^21,22^ show states both with the π-helix and with an α-helix in this region (Supplementary Figure 12B, C). In contrast, the TRPV2 structures presented in this paper as well as those of the minimal TRPV2 solved by both cryo-EM^20^ and x-ray crystallography^31^ only show α-helices for the whole S6 helix. It’s unclear whether the presence of the π-helix would hinder or aid CBD binding and CBD-based opening of the channel.

To our surprise, the presence of various lipids and PI(4,5)P_2_ during nanodisc reconstitution did not reveal the binding site for PI(4,5)P_2_ and did not allow us to determine the proposed^18,24^ role PI(4,5)P_2_ plays in channel gating. Nevertheless, our new TRPV2 structures confirmed that the absence of obvious lipid density in the vanilloid pocket for any TRPV2 cryo-EM structure so far^18–20^ is a characteristic feature of the channel. Further investigation needs to be done on other endogenous membrane lipids that may modulate the channel.

Overall, our structural studies have revealed a novel drug binding site in TRPV channels and provided molecular insights into TRPV2 modulation by CBD that could be used to guide therapeutic design to treat glioblastoma multiforme and other TRPV2 channel associated pathophysiological process.

## METHODS

### Protein expression and purification

The detergent solubilized full-length rat TRPV2 was expressed and purified as described previously^20^. Briefly, rat TRPV2 expressing plasma membranes from *S. cerevisiae* were solubilized in a buffer containing 20 mM HEPES pH 8.0, 150 mM NaCl, 5% glycerol, 0.087% LMNG (Anatrace), 2 mM Tris (2-carboxyethyl) phosphinehydrochloride (TCEP), 1 mM Phenylmethylsulfonyl fluoride (PMSF) and supplemented with protease inhibitor cocktail tablet mini (Roche), for 1 hour. Detergent insoluble material was removed by ultra-centrifugation at 100,000 × *g* and the protein was purified from the supernatant by affinity chromatography by binding with 1D4 antibody coupled CnBr-activated Sepharose beads. A column was packed with the Sepharose beads and washed with Wash Buffer (20 mM HEPES pH 8.0, 150 mM NaCl, 2 mM TCEP) containing 0.006% DMNG (Anatrace). TRPV2 was eluted with Wash Buffer containing 0.006% DMNG and 3 mg/ml 1D4 peptide (GenScript USA) and subjected to size-exclusion chromatography using a Superose 6 column (GE Healthcare) with Wash Buffer containing 0.006% DMNG. Fractions containing TRPV2 protein were concentrated to ~3 mg/mL and used for vitrification.

The nanodisc reconstituted full-length rat TRPV2 was expressed and purified as previously published^16^, with minor modifications. The membranes expressing rat TRPV2 were solubilized in 20 mM HEPES pH 8.0, 150 mM NaCl, 5% glycerol, 0.087% LMNG, 2 mM TCEP, and 1 mM PMSF for 1 hour. Insoluble material was removed via ultra-centrifugation at 100,000 × *g* and the solubilized TRPV2 was purified by binding to 1D4 antibody coupled CnBr-activated Sepharose beads. The beads were washed with Wash Buffer containing 0.006% DMNG and the protein was eluted with Wash Buffer containing 0.006% DMNG and 3 mg/ml 1D4 peptide. The protein was incubated with 100 μM cannabidiol (CBD, Cayman Chemical Company) and 400 μM diC8 PI(4,5)P_2_ (Echelon Biosciences) for 1 hour. The liganded protein was then reconstituted into nanodiscs in a 1:1:200 ratio of TRPV2:MSP2N2:soy polar lipids (Avanti). MSP2N2 was expressed in BL21 (DE3) cells and purified via affinity chromatography as previously described^16^. Lipids were dried under nitrogen flow for 3 hours prior to reconstitution and resuspended in Wash Buffer containing DMNG in a 1:2.5 ratio (soy polar lipids:DMNG). The nanodisc reconstitution mixture was incubated at 4°C for 30 mins. Bio-Beads (Bio-Beads SM-2 Absorbent Media, Bio-Rad) were added to the reconstitution mixture for 1 hour then fresh Bio-Beads were added and the mixture was allowed to incubate overnight. Nanodisc reconstituted TRPV2 was further purified using size-exclusion chromatography (Superose 6, GE Healthcare) in Wash Buffer. Protein eluted from the column was concentrated to 1.9 mg/mL for use in vitrification.

### Cryo-EM data collection

For the detergent solubilized TRPV2, prior to preparing cryo-EM grids, purified TRPV2 was incubated with 30 μM CBD for 30 min. Fluorinated Fos-choline 8 was added to sample to a final concentration of 3 mM just before blotting. This sample was double blotted (3.5 μl per blot) onto 200 mesh Quantifoil 1.2/1.3 grids (Quantifoil Micro Tools) at 4°C and 100% humidity and plunge frozen in liquid ethane cooled to the temperature of liquid nitrogen (FEI Vitrobot). Cryo-EM images were collected using a 300kV FEI Titan Krios microscope equipped with a Gatan K2 Summit direct detector camera. Images were recorded using super resolution counting mode following the established protocol. Specifically, 38 frame movies were collected at 18,000x magnification with a physical pixel size of 0.69Å/pix and a dose rate of 6.85 e/pix/s. Total exposure time was 11.4 seconds with one frame recorded every 0.3 seconds using the automated imaging software, Leginon^32^. Defocus values of the images ranged from −1.4 to - 3.0 μm.

For the nanodisc reconstituted TRPV2, prior to preparing cryo-EM grids, purified TRPV2 was incubated with 100 μM CBD and 400 μM diC8 PI(4,5)P_2_ for 30 mins. Fluorinated Fos-choline 8 was added to the sample to a final concentration of 3 mM just before blotting. This sample was double blotted (3.5 μl per blot) onto 200 mesh Quantifoil 1.2/1.3 grids (Quantifoil Micro Tools) at 4°C and 100% humidity and plunge frozen in liquid ethane cooled to the temperature of liquid nitrogen (FEI Vitrobot). Cryo-EM images were collected using a 300kV FEI Titan Krios microscope equipped with a Gatan K2 Summit direct detector camera. 50 frame movies were collected at with a dose rate of 8.0 e/pix/s with 5 frames per second and a super resolution pixel size of 0.535 Å/pix. Defocus values of the images range from −0.8 to −3.0 μm.

### Image processing

For the detergent solubilized TRPV2 in the presence of CBD, the movie frames were aligned and binned using MotionCor2^29^ to compensate for beam-induced motion. All subsequent data processing was conducted in RELION 2.1^27,33,34^. Defocus values of the motion corrected micrographs were estimated using Gctf^30^. Initially, ~2,000 particles were manually picked from 3,779 micrographs and sorted into 2D classes to generate templates for auto-picking. Auto-picking with a loose threshold to ensure maximum picking resulted in ~821,000 auto-picked particles. These were then subjected to 2D classification to remove suboptimal particles and false positive hits. The best ~258,000 particles were then auto-refined using the 3D auto-refinement option without masking or applied symmetry. The initial model for this refinement was created from the density map of the previously published full length TRPV2^19^ (EMB-6580) filtered to 60 Å. The ~258,000 particles were then 3D classified into 8 classes with masking, but without aligning angles to the particles. This initial model used for this classification was the reconstruction of the ~258,000 particles described above. The mask used in this classification was created from the initial refinement of the ~258,000 particles adjusted to a threshold of 0.005, filtered to 15Å, extended by 5 pix and a soft edge of 5 pix was added. One distinct class with four-fold symmetry contained 49,496 particles and was refined with C4 symmetry using the mask and initial model described above. The map from this refinement was used as an initial model for further classification into 3 classes without applying angles. This classification was masked using the map refined from the 49,496 particles adjusted to a threshold of 0.005, filtered to 5Å, extended by 5 pix with a soft edge of 5 pix. The most stable class contained 26,322 particles and was refined with the same initial model and mask to produce a 4.6 Å map with C4 symmetry. Postprocessing yielded a 4.3Å resolution map as estimated by Rmeasure^35^. Local resolutions were estimated using the RESMAP software^36^.

For the nanodisc reconstituted TRPV2 in the presence of CBD and diC8 PI(4,5)P_2_, the movie frames were aligned and binned using MotionCor2^29^ to compensate for beam-induced motion. All subsequent data processing was conducted in RELION 3.0^28,33,34^. Defocus values of the motion corrected micrographs were estimated using Gctf^30^. Initially, ~3,000 particles were manually picked and sorted into 2D classes to generate templates for auto-picking. Auto-picking of 7,061 micrographs resulted in ~685,000 auto-picked particles. These were then subjected to 2D classification to remove suboptimal particles and false positive hits. The best ~348,000 particles were refined without applied symmetry. The initial model for this refinement was created from the density map of the previously published full length TRPV2^19^ (EMB-6580) filtered to 60 Å. These ~348,000 particles were then subjected to 3D classification into 8 classes with angular sampling and no applied symmetry, using a mask made from the initial 3D refinement. The mask was made by lowpass filtering the map to 5 Å, with a threshold of 0.004, and the map extended by 10 pix with a soft edge of 10 pix. The best class had ~68,000 particles and was refined with C4 symmetry in a mask created from the 3D class. This mask was made by lowpass filtering the 3D class to 5 Å, with a threshold of 0.006, and the map extended by 10 pix with a soft edge of 10 pix. These particles were then subjected to CTF refinement and Bayesian polishing. The particles were then 3D refined again using a mask made from the map of the ~68,000 unpolished particles, lowpass filtered to 10 Å, at a threshold of 0.005, extended 7 pix with a soft edge of 7 pix. These particles were then subjected to 3D classification into 3 classes, with no alignments. The best class contained ~25,000 particles and was refined with C4 symmetry using the same mask as the previous refinement. Postprocessing yielded a map at 3.5 Å. Local resolutions were estimated using the RESMAP software^36^.

The apo TRPV2 in detergent movie frames were aligned and binned using MotionCor2^29^ to compensate for beam-induced motion. All subsequent data processing was conducted in RELION 3.0^28,33,34^. Defocus values of the motion corrected micrographs were estimated using Gctf^30^. Approximately 26,000 particles were autopicked from a subset of 10 micrographs using template-free Laplacian-of-Gaussian autopicking. These particles were then subjected to 2D classification to create templates for autopicking. These templates were used to autopick from the full 988 micrograph dataset, yielding ~420,000 particles. These particles were then subject to 2D classification. The best ~278,000 particles were then auto-refined using the 3D auto-refinement option without masking or applied symmetry. The initial model for this refinement was created from the density map of the previously published full length TRPV2^19^ (EMB-6580) filtered to 60 Å. The ~278,000 particles were then 3D classified into 8 classes with masking, but without aligning angles to the particles. The initial model used for this classification was the reconstruction of the ~278,000 particles described above. The mask used in this classification was created from the initial refinement of the ~278,000 particles adjusted to a threshold of 0.005, filtered to 15Å, extended by 5 pix and a with a soft edge of 5 pix added. The best class from this classification contained ~51,000 particles which were refined with C4 applied symmetry with the same initial model and mask described above. These particles were then subjected to CTF refinement and Bayesian polishing. The polished ~51,000 particles were refined with applied C4 symmetry, with a mask made from the map of the previously refined ~51,000 particles at threshold 0.005, extended by 5 pix with a soft edge of 5 pix and using that same previous map as the reference model. Post-processing yielded a map at 4.2 Å. Local resolutions were estimated using the RESMAP software^36^.

### Model building

The previously determined full-length rat TRPV2 structure (PDB: 5HI9) was employed as our initial starting model and docked into the CBD-bound TRPV2 in nanodiscs and apo TRPV2 maps. This model was then manually adjusted to each map in COOT^37^ and then refined using phenix.real_space_refine from the PHENIX software package^38^ with four-fold NCS constraints. The models were subjected to iterative rounds of manual model fitting followed by real-space refinement and sidechains with insufficient density were removed. Due to the lower quality of the CBD-bound TRPV2 in detergent map, we started from the already refined CBD-bound TRPV2 in nanodiscs model. This model was adjusted manually and again refined using multiple rounds of manual model building in COOT and phenix.real_space_refine. The disconnected 16 residue loop between the ARDs and the β-sheet region in the CBD-bound TRPV2 in detergent model was built as alanines and arbitrarily assigned as residues 1-16.

The final models were randomized by 0.5 Å in PHENIX^38^ and refined against each half map. These models were converted into volumes in Chimera^39^ and EMAN2^40^ used to generate FSC curves between these models and their half maps as well as between each final model and their summed maps. HOLE was used to generate the pore radii^41^. Pymol and Chimera^39^ were used to align models and maps and to make figures.

### HEK293 cell culture, mutagenesis and transfection

Human Embryonic Kidney 293 (HEK293) cells were purchased from American Type Culture Collection (ATCC), Manassas, VA, (catalogue # CRL-1573), RRID:CVCL_0045. The cells were maintained in minimal essential medium (MEM) (Life Technologies, Carlsbad, CA, USA) supplemented with 10% (v/v) fetal bovine serum (FBS), 100 IU/ml penicillin and 100 μg/ml streptomycin (37°C in 5% CO_2_). The cells were transiently transfected with cDNA encoding the rat TRPV2 and its mutants using the Effectene reagent (Qiagen) 48-72 hours before experiments. Point mutations were introduced using the QuickChange Mutagenesis Kit (Agilent).

### Ca^2+^ imaging

Ca^2+^ imaging measurements were performed with an Olympus IX-51 inverted microscope equipped with a DeltaRAM excitation light source (Photon Technology International, Horiba), as described earlier^42^. Briefly, HEK cells were loaded with 1 μM fura-2 AM (Invitrogen) at 37°C for 40-50 min, fluorescence images were collected with a Roper Cool-Snap digital CCD camera at 510 nm emission wavelength, excitation light was provided by a DeltaRAM light source alternating between 340 and 380 nm. Measurements were conducted in an extracellular solution containing (in mM) 137 NaCl, 5 KCl, 1 MgCl_2_, 2 CaCl_2_, 10 HEPES and 10 glucose, pH 7.4. Data analysis was performed using the Image Master software (PTI).

## Acknowledgments

We thank David Lodowski at Case Western Reserve University for help in the early stage of the project. We thank Sudha Chakrapani at Case Western Reserve University for assistance with MSP2N2 expression and purification. We thank Sabine Baxter for assistance with hybridoma and cell culture at the University of Pennsylvania Perelman School of Medicine Cell Center Services Facility. We thank Sudheer Molugu for assistance with microscopes operation and acknowledge the use of instruments at the Electron Microscopy Resource Lab at the University of Pennsylvania Perelman School of Medicine. We also acknowledge the use of instruments at the Electron Imaging Center for NanoMachines supported by NIH (1S10RR23057 and 1S10OD018111), NSF (DBI-1338135) and CNSI at UCLA. This work was supported by grants from the National Institute of Health (R01GM103899 and R01GM129357 to V.Y.M.-B., U24 GM116792 to Z.H.Z and V.Y.M.-B, R01NS055159 and R01GM093290 to T.R.).

## Authors Contributions

R.A.P. performed protein purifications, cryo-EM sample preparation, all cryo-EM data analysis and interpretation, and built and refined all atomic models; A.S. conducted initial cryo-EM studies, including protein purification, sample preparation, and initial cryo-EM data analysis; Y.L. performed cryo-EM data collection at Pfizer; T.E.T.H. aided in protein purification and cryo-EM sample preparation; S.Z. and Y.Y. performed Ca^2+^ imaging experiments; K.W.H performed data collection at UCLA; Z.H.Z. supervised cryo-EM data collection at UCLA; T.R. supervised Ca^2+^ imaging experiments; S.H. supervised cryo-EM data collection at Pfizer; V.Y.M-B. designed and supervised the execution of all experiments in this manuscript; R.A.P., A.S., T.E.T.H. and V.Y.M-B. drafted the manuscript; all authors contributed to and reviewed the final manuscript.

## Data availability

The cryo-EM density maps and the atomic coordinates of the apo and both CBD-bound full-length TRPV2 channels are deposited into the Electron Microscopy Data Bank and Protein Data Bank under accession codes EMD-0461 and PDB 6NNK (apo state), EMD-0462 and PDB 6NNL (CBD-bound in detergent) and EMD-0463 and PDB 6NNM (CBD-bound in nanodiscs).

## REFERENCES

1 Venkatachalam, K., Luo, J. & Montell, C. Evolutionarily conserved, multitasking TRP channels: lessons from worms and flies. Handb Exp Pharmacol 223, 937–962, doi:10.1007/978-3-319-05161-1_9 (2014).

2 Venkatachalam, K. & Montell, C. TRP channels. Annu Rev Biochem 76, 387–417, doi:10.1146/annurev.biochem.75.103004.142819 (2007).

3 Park, U. et al. TRP vanilloid 2 knock-out mice are susceptible to perinatal lethality but display normal thermal and mechanical nociception. J Neurosci 31, 11425–11436, doi:10.1523/JNEUROSCI.1384-09.2011 (2011).

4 Qin, N. et al. TRPV2 is activated by cannabidiol and mediates CGRP release in cultured rat dorsal root ganglion neurons. J Neurosci 28, 6231–6238, doi:10.1523/JNEUROSCI.0504-08.2008 (2008).

5 Liberati, S. et al. Loss of TRPV2 Homeostatic Control of Cell Proliferation Drives Tumor Progression. Cells 3, 112–128, doi:10.3390/cells3010112 (2014).

6 Nabissi, M. et al. TRPV2 channel negatively controls glioma cell proliferation and resistance to Fas-induced apoptosis in ERK-dependent manner. Carcinogenesis 31, 794–803, doi:10.1093/carcin/bgq019 (2010).

7 Nabissi, M. et al. Cannabidiol stimulates Aml-1a-dependent glial differentiation and inhibits glioma stem-like cells proliferation by inducing autophagy in a TRPV2-dependent manner. Int J Cancer 137, 1855–1869, doi:10.1002/ijc.29573 (2015).

8 Nabissi, M., Morelli, M. B., Santoni, M. & Santoni, G. Triggering of the TRPV2 channel by cannabidiol sensitizes glioblastoma cells to cytotoxic chemotherapeutic agents. Carcinogenesis 34, 48–57, doi:10.1093/carcin/bgs328 (2013).

9 Peralvarez-Marin, A., Donate-Macian, P. & Gaudet, R. What do we know about the transient receptor potential vanilloid 2 (TRPV2) ion channel? FEBS J 280, 5471–5487, doi:10.1111/febs.12302 (2013).

10 De Petrocellis, L. et al. Cannabinoid actions at TRPV channels: effects on TRPV3 and TRPV4 and their potential relevance to gastrointestinal inflammation. Acta Physiol (Oxf) 204, 255–266, doi:10.1111/j.1748-1716.2011.02338.x (2012).

11 Smart, D. et al. The endogenous lipid anandamide is a full agonist at the human vanilloid receptor (hVR1). Br J Pharmacol 129, 227–230, doi:10.1038/sj.bjp.0703050 (2000).

12 De Petrocellis, L. et al. Plant-derived cannabinoids modulate the activity of transient receptor potential channels of ankyrin type-1 and melastatin type-8. J Pharmacol Exp Ther 325, 1007–1015, doi:10.1124/jpet.107.134809 (2008).

13 Madej, M. G. & Ziegler, C. M. Dawning of a new era in TRP channel structural biology by cryo-electron microscopy. Pflugers Arch 470, 213–225, doi:10.1007/s00424-018-2107-2 (2018).

14 Rohacs, T. Phosphoinositide regulation of TRP channels. Handb Exp Pharmacol 223, 1143–1176, doi:10.1007/978-3-319-05161-1_18 (2014).

15 Gao, Y., Cao, E., Julius, D. & Cheng, Y. TRPV1 structures in nanodiscs reveal mechanisms of ligand and lipid action. Nature 534, 347–351, doi:10.1038/nature17964 (2016).

16 Hughes, T. E. T. et al. Structural basis of TRPV5 channel inhibition by econazole revealed by cryo-EM. Nat Struct Mol Biol 25, 53–60, doi:10.1038/s41594-017-0009-1 (2018).

17 McGoldrick, L. L. et al. Opening of the human epithelial calcium channel TRPV6. Nature 553, 233–237, doi:10.1038/nature25182 (2018).

18 Dosey, T. L. et al. Structures of TRPV2 in distinct conformations provide insight into role of the pore turret. Nat Struct Mol Biol 26, 40–49, doi:10.1038/s41594-018-0168-8 (2019).

19 Huynh, K. W. et al. Structure of the full-length TRPV2 channel by cryo-EM. Nat Commun 7, 11130, doi:10.1038/ncomms11130 (2016).

20 Zubcevic, L. et al. Cryo-electron microscopy structure of the TRPV2 ion channel. Nat Struct Mol Biol 23, 180–186, doi:10.1038/nsmb.3159 (2016).

21 Singh, A. K., McGoldrick, L. L. & Sobolevsky, A. I. Structure and gating mechanism of the transient receptor potential channel TRPV3. Nat Struct Mol Biol 25, 805–813, doi:10.1038/s41594-018-0108-7 (2018).

22 Zubcevic, L. et al. Conformational ensemble of the human TRPV3 ion channel. Nat Commun 9, 4773, doi:10.1038/s41467-018-07117-w (2018).

23 Deng, Z. et al. Cryo-EM and X-ray structures of TRPV4 reveal insight into ion permeation and gating mechanisms. Nat Struct Mol Biol 25, 252–260, doi:10.1038/s41594-018-0037-5 (2018).

24 Mercado, J., Gordon-Shaag, A., Zagotta, W. N. & Gordon, S. E. Ca2+-dependent desensitization of TRPV2 channels is mediated by hydrolysis of phosphatidylinositol 4,5-bisphosphate. J Neurosci 30, 13338–13347, doi:10.1523/JNEUROSCI.2108-10.2010 (2010).

25 Velisetty, P. et al. A molecular determinant of phosphoinositide affinity in mammalian TRPV channels. Sci Rep 6, 27652, doi:10.1038/srep27652 (2016).

26 Cao, E., Liao, M., Cheng, Y. & Julius, D. TRPV1 structures in distinct conformations reveal activation mechanisms. Nature 504, 113–118, doi:10.1038/nature12823 (2013).

27 Kimanius, D., Forsberg, B. O., Scheres, S. H. & Lindahl, E. Accelerated cryo-EM structure determination with parallelisation using GPUs in RELION-2. Elife 5, doi:10.7554/eLife.18722 (2016).

28 Zivanov, J. et al. New tools for automated high-resolution cryo-EM structure determination in RELION-3. Elife 7, doi:10.7554/eLife.42166 (2018).

29 Zheng, S. Q. et al. MotionCor2: anisotropic correction of beam-induced motion for improved cryo-electron microscopy. Nat Methods 14, 331–332, doi:10.1038/nmeth.4193 (2017).

30 Zhang, K. Gctf: Real-time CTF determination and correction. J Struct Biol 193, 1–12, doi:10.1016/j.jsb.2015.11.003 (2016).

31 Zubcevic, L., Le, S., Yang, H. & Lee, S. Y. Conformational plasticity in the selectivity filter of the TRPV2 ion channel. Nat Struct Mol Biol 25, 405–415, doi:10.1038/s41594-018-0059-z (2018).

32 Potter, C. S. et al. Leginon: a system for fully automated acquisition of 1000 electron micrographs a day. Ultramicroscopy 77, 153–161 (1999).

33 Scheres, S. H. Processing of Structurally Heterogeneous Cryo-EM Data in RELION. Methods Enzymol 579, 125–157, doi:10.1016/bs.mie.2016.04.012 (2016).

34 Scheres, S. H. RELION: implementation of a Bayesian approach to cryo-EM structure determination. J Struct Biol 180, 519–530, doi:10.1016/j.jsb.2012.09.006 (2012).

35 Sousa, D. & Grigorieff, N. Ab initio resolution measurement for single particle structures. J Struct Biol 157, 201–210, doi:10.1016/j.jsb.2006.08.003 (2007).

36 Kucukelbir, A., Sigworth, F. J. & Tagare, H. D. Quantifying the local resolution of cryo-EM density maps. Nat Methods 11, 63–65, doi:10.1038/nmeth.2727 (2014).

37 Emsley, P. & Cowtan, K. Coot: model-building tools for molecular graphics. Acta Crystallogr D Biol Crystallogr 60, 2126–2132, doi:10.1107/S0907444904019158 (2004).

38 Adams, P. D. et al. PHENIX: building new software for automated crystallographic structure determination. Acta Crystallogr D Biol Crystallogr 58, 1948–1954 (2002).

39 Pettersen, E. F. et al. UCSF Chimera--a visualization system for exploratory research and analysis. J Comput Chem 25, 1605–1612, doi:10.1002/jcc.20084 (2004).

40 Ludtke, S. J., Baldwin, P. R. & Chiu, W. EMAN: semiautomated software for high-resolution single-particle reconstructions. J Struct Biol 128, 82–97, doi:10.1006/jsbi.1999.4174 (1999).

41 Smart, O. S., Neduvelil, J. G., Wang, X., Wallace, B. A. & Sansom, M. S. HOLE: a program for the analysis of the pore dimensions of ion channel structural models. J Mol Graph 14, 354–360, 376 (1996).

42 Lukacs, V. et al. Distinctive changes in plasma membrane phosphoinositides underlie differential regulation of TRPV1 in nociceptive neurons. J Neurosci 33, 11451–11463, doi:10.1523/JNEUROSCI.5637-12.2013 (2013).

