## Supplementary Material for "Molecular mechanism of TRPV2 channel modulation by cannabidiol"

**Supplementary Figure 1. EM summary of CBD-bound TRPV2 in detergent.** (**A**) Representative micrograph and 2D classes of cryo-EM data. (**B**) Map FSC curves and model validation curves. (**C**) Angular distribution for final 3D reconstruction. Tall red cylinders indicate a large number of particles aligned at a position and short blue cylinders indicate fewer particles. (**D**) Top, bottom and side view of CBD-bound TRPV2 in detergent colored based on local resolution as calculated via RESMAP. Blue, white and red regions indicate resolutions of 3.5Å, 4.5Å and 5.5Å, respectively.

**Supplementary Figure 2. CBD-bound TRPV2 in detergent.** (**A**) Side and top views of the structure of CBD-bound TRPV2 in detergent solved to 4.3Å. A single monomer is colored in green and the others are shown in grey. The densities attributed to lipids are shown in khaki. (**B**) Atomic model of CBD-bound TRPV2 in detergent. A single monomer is colored in green and the others are shown in grey. (**C**) Schematic overview of TRPV2 domains.

**Supplementary Figure 3. Data quality of the CBD-bound TRPV2 in detergent structure.** Selected helices of the CBD-bound TRPV2 in detergent model (green) overlaid with the corresponding cryo-EM density map in grey mesh. Side chains are shown as sticks and atoms are colored by element.

**Supplementary Figure 4. EM summary of apo TRPV2 in detergent.** (**A**) Representative micrograph and 2D classes of cryo-EM data. (**B**) Map FSC curves and model validation curves. (**C**) Angular distribution for final 3D reconstruction. Tall red cylinders indicate a large number of particles aligned at a position and short blue cylinders indicate fewer particles. (**D**) Top, bottom and side view of apo TRPV2 colored based on local resolution as calculated via RESMAP. Blue, white and red regions indicate resolutions of 3.5Å, 4.5Å and 5.5Å, respectively.

**Supplementary Figure 5. Apo TRPV2 in detergent.** (**A**) Side and top views of the structure of apo TRPV2 in detergent solved to 4.1Å. A single monomer is colored in salmon and the others are shown in grey. The densities attributed to lipids are shown in khaki. (**B**) Atomic model of apo TRPV2 in detergent. A single monomer is colored in salmon and the others are shown in grey.

**Supplementary Figure 6. Data quality of the apo TRPV2 in detergent structure.** Selected helices of the apo TRPV2 model (salmon) overlaid with the corresponding cryo-EM density map in grey mesh. Side chains are shown as sticks and atoms are colored by element.

**Supplementary Figure 7. EM summary of CBD-bound TRPV2 in PI(4,5)$P_2$ enriched nanodiscs**. (**A**) Representative micrograph and 2D classes of cryo-EM data. (**B**) Map FSC curves and model validation curves. (**C**) Angular distribution for final 3D reconstruction. Tall red cylinders indicate a large number of particles aligned at a position and short blue cylinders indicate fewer particles. (**D**) Top, bottom and side view of nanodisc reconstituted CBD-bound TRPV2 in the presence of PI(4,5)$P_2$ colored based on local resolution as calculated via RESMAP. Blue, white and red regions indicate resolutions of 3.5Å, 4.5Å and 5.5Å, respectively.

**Supplementary Figure 8. CBD-bound TRPV2 in PI(4,5)$P_2$ enriched nanodiscs**. (**A**) Side and top views of the structure of CBD-bound TRPV2 in PI(4,5)$P_2$ enriched nanodiscs solved to 4.1Å. A single monomer is colored in blue and the others are shown in grey. The densities attributed to bound CBD and lipids are shown in pink and khaki, respectively. (**B**) Atomic model of CBD-bound TRPV2 in PI(4,5)$P_2$ enriched nanodiscs. A single monomer is colored in blue and the others are shown in grey. Bound CBD is shown as pink sticks.

**Supplementary Figure 9. Data quality of the CBD-bound TRPV2 in nanodiscs structure.** Selected helices of the CBD-bound TRPV2 in the presence of PI(4,5)$P_2$ model (blue) overlaid with the corresponding cryo-EM density map in grey mesh. Side chains are shown as sticks and atoms are colored by element.

**Supplementary Figure 10. CBD binding pocket.** (**A**) Chemical diagram of molecular cannabidiol (CBD). (**B**) The CBD binding pocket of CBD-bound TRPV2 in PI(4,5)$P_2$ enriched nanodiscs. The electron density map of the S5 and S6 helices of TRPV2 are shown as blue and grey surfaces, respectively. The density attributed to bound CBD is shown as pink mesh and the chemical structure of CBD is shown as pink sticks. (**C**) Model representation of the CBD binding pocket. The model of the S5 and S6 helices of CBD-bound TRPV2 in nanodiscs are shown as blue and grey cartoons, respectively. This model is overlaid with the S5 and S6 helices of apo TRPV2 in detergent which are shown as pink and grey cartoons, respectively. Residues of interest are labeled and represented as sticks.

**Supplementary Figure 11. CBD cryo-EM density.** TRPV2 models overlaid with the sharpened density map and the two corresponding half maps for CBD-bound TRPV2 in PI(4,5)$P_2$ enriched nanodiscs (top, blue), CBD-bound TRPV2 in detergent (middle, green) and apo TRPV2 in detergent (bottom, pink). Cryo-EM densities shown as grey mesh and side chains of interest are shown as sticks. Bound CBD (top) is shown as pink sticks.

**Supplementary Figure 12. Tyrosine 525 comparison.** Area of S4-S5 linker for (A) CBD-bound TRPV2 in nanodiscs, (B) apo human TRPV3 (PDB 6MHO) and (C) 2-APB sensitized human TRPV3 (PDB 6MHS). Residues of interest are labeled and shown as sticks.

**Supplementary Figure 13. The L541F-L631F double mutation enhances CBD responses of TRPV2.** $Ca^{2+}$ imaging experiments on Fura-2 loaded HEK293 cells transfected either with the wild type rat TRPV2 or the L541F-L631F double mutant were performed as described in the methods section. (A) $Ca^{2+}$ imaging traces show mean +/- S.E.M, n=861 cells from 5 coverslips for wild type TRPV2 and n=961 cells for 6 coverslips for L541- L631F. The applications of 20 μM CBD, 50 μM CBD and 100 μM 2-APB are indicated by the horizontal lines. (B) Summary data; the responses to 20 μM or 50 μM CBD were normalized to those evoked by 100 μM 2APB. ***$p<0.001$

**Supplementary Figure 14. Lipid binding pocket.** The lipid binding pocket located between the S1, S2 and TRP helix of (**A**) apo TRPV2 in detergent (pink) (**B**) CBD-bound TRPV2 in detergent, (C) CBD-bound TRPV2 in PI(4,5)$P_2$ enriched nanodiscs, and (D) apo TRPV1 in nanodiscs (PDB 5IRZ). The models for the helices that constitute the pockets are shown as ribbons and overlaid with their respective cryo-EM densities shown as grey surfaces. Arrows indicate the location of non-protein densities in these pockets.

**Supplementary Figure 15. Map quality and pore comparisons of WT TRPV2.** Ribbon representation of the S4-S5 linker, S5 and S6 helices overlaid with their respective cryo-EM maps as well as the pore dimer cartoons for the reprocessed apo TRPV2 in detergent (salmon, left), TRPV2 in the open state (green, center) and the previously published apo TRPV2 (light pink, right). Cryo-EM maps are shown as grey mesh. Pore diameter measurements are from the carbon backbone of each model.

**Supplementary Figure 16. Residue conservation in the CBD-binding pocket**. Sequence alignment of select TRPV family channels of the S5-S6 region. Residues involved with CBD-binding are highlighted in yellow.

### Supplementary Figure 1

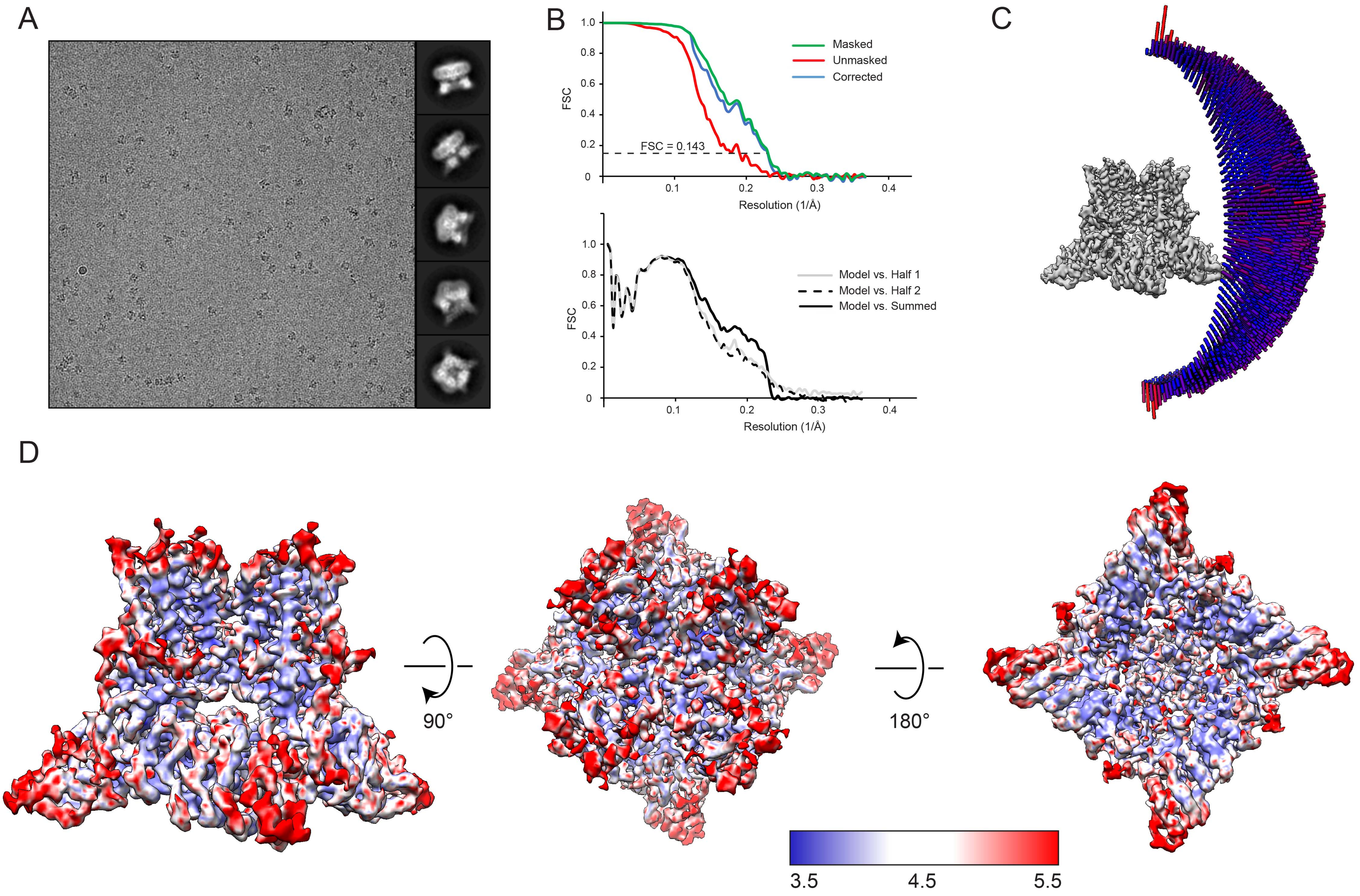

Supplementary Figure 2

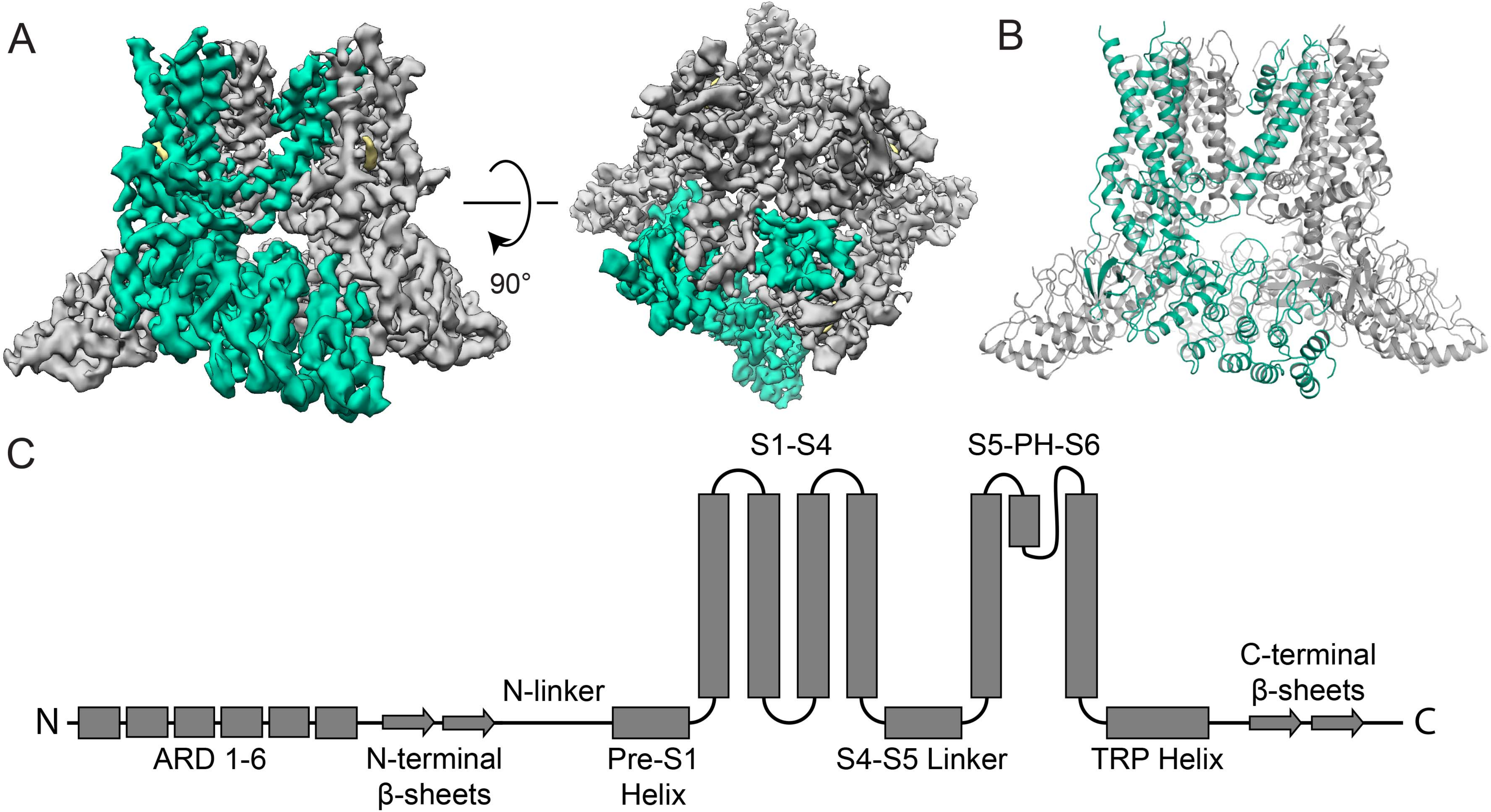

### Supplementary Figure 3

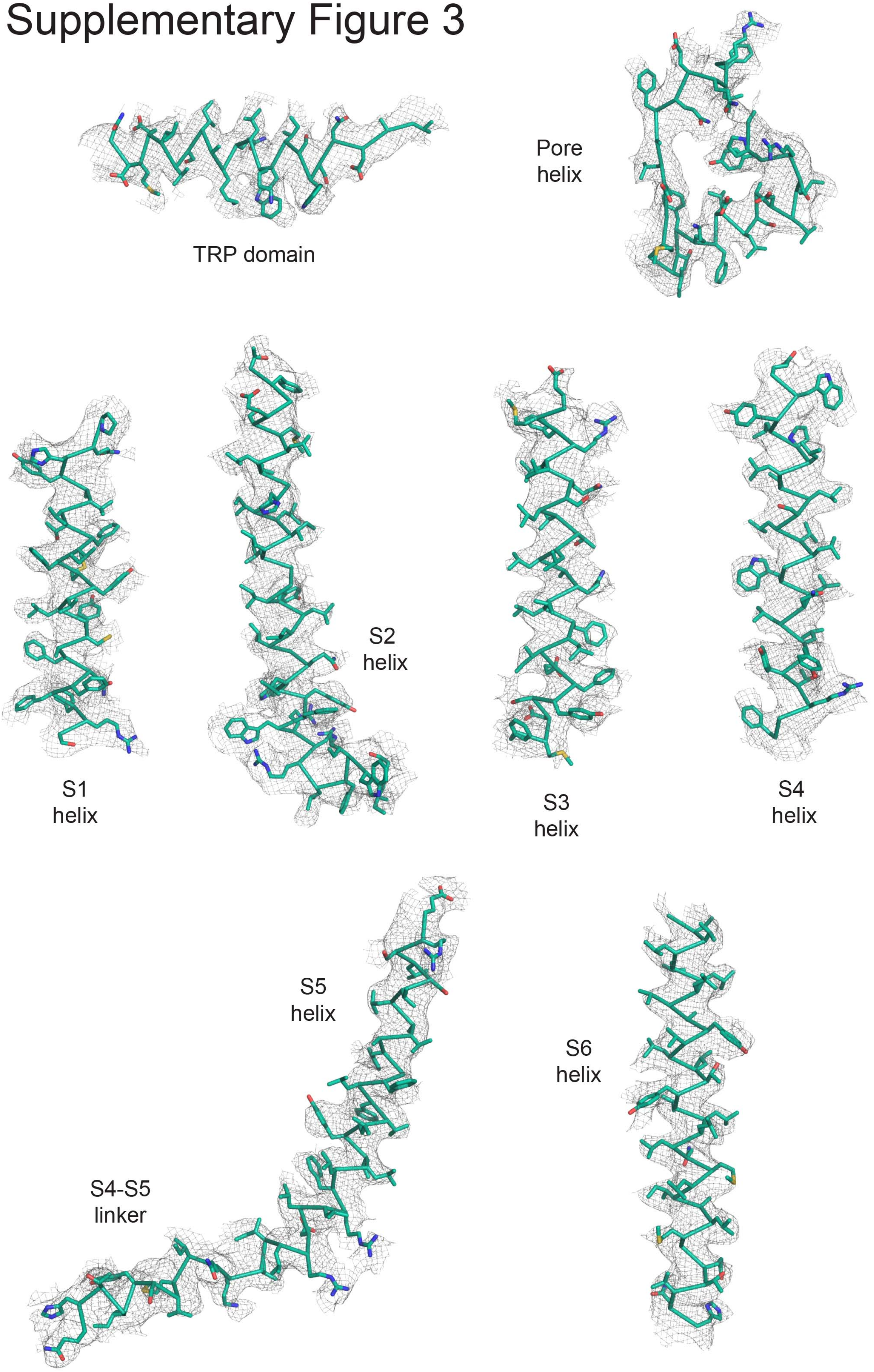

Supplementary Figure 4

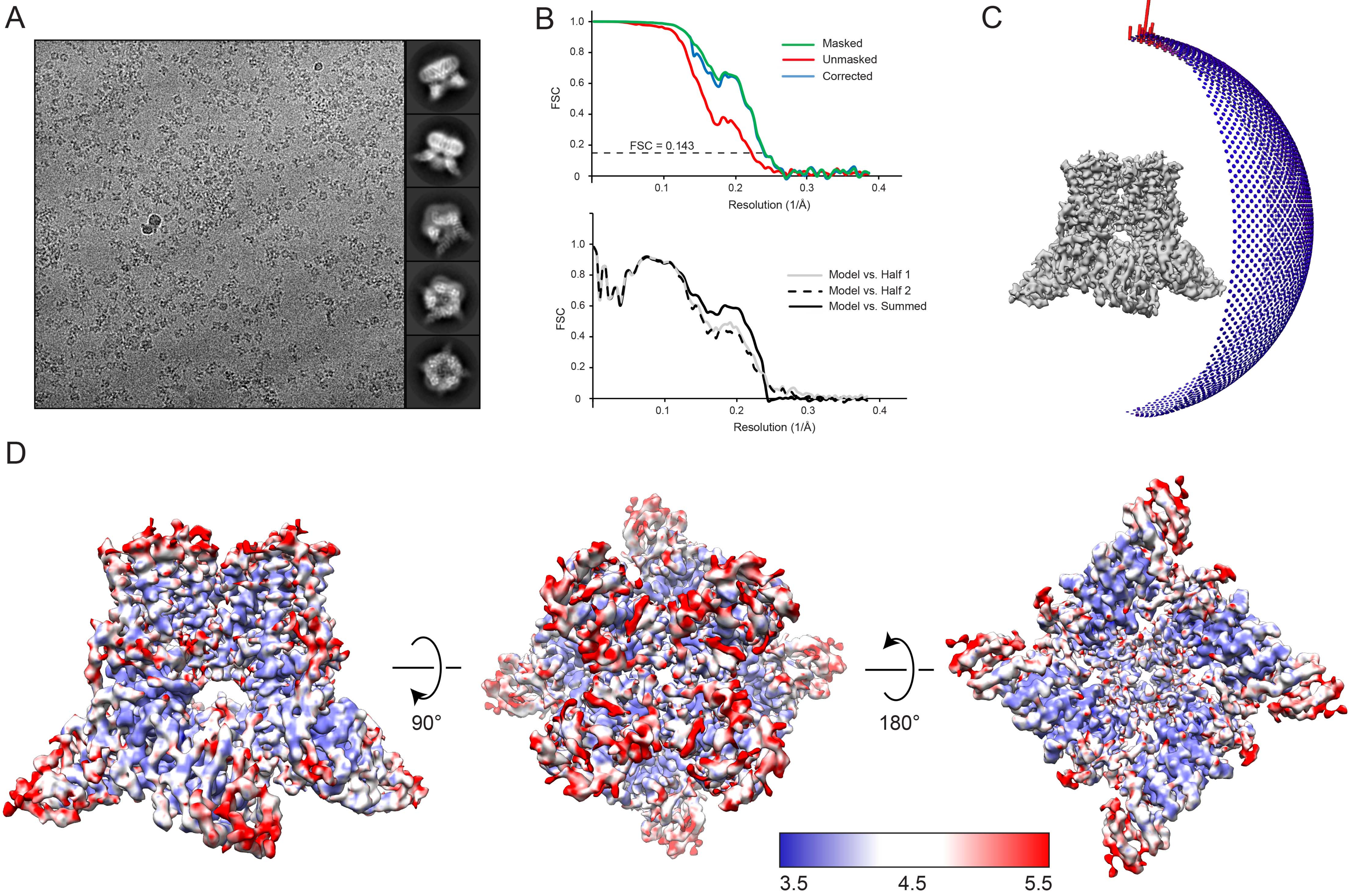

Supplementary Figure 5

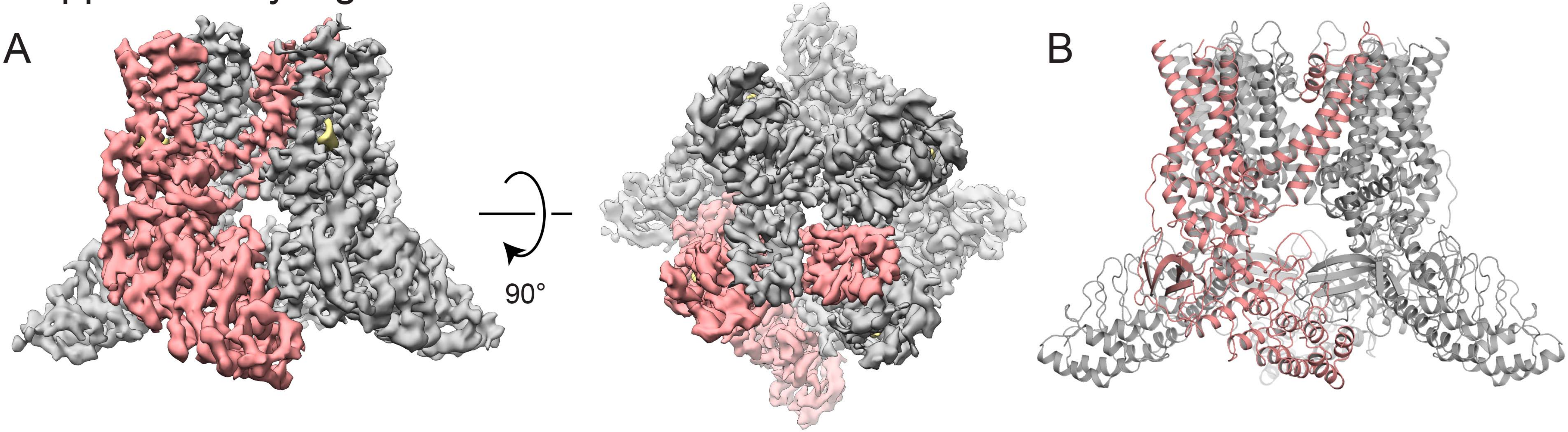

### Supplementary Figure 6

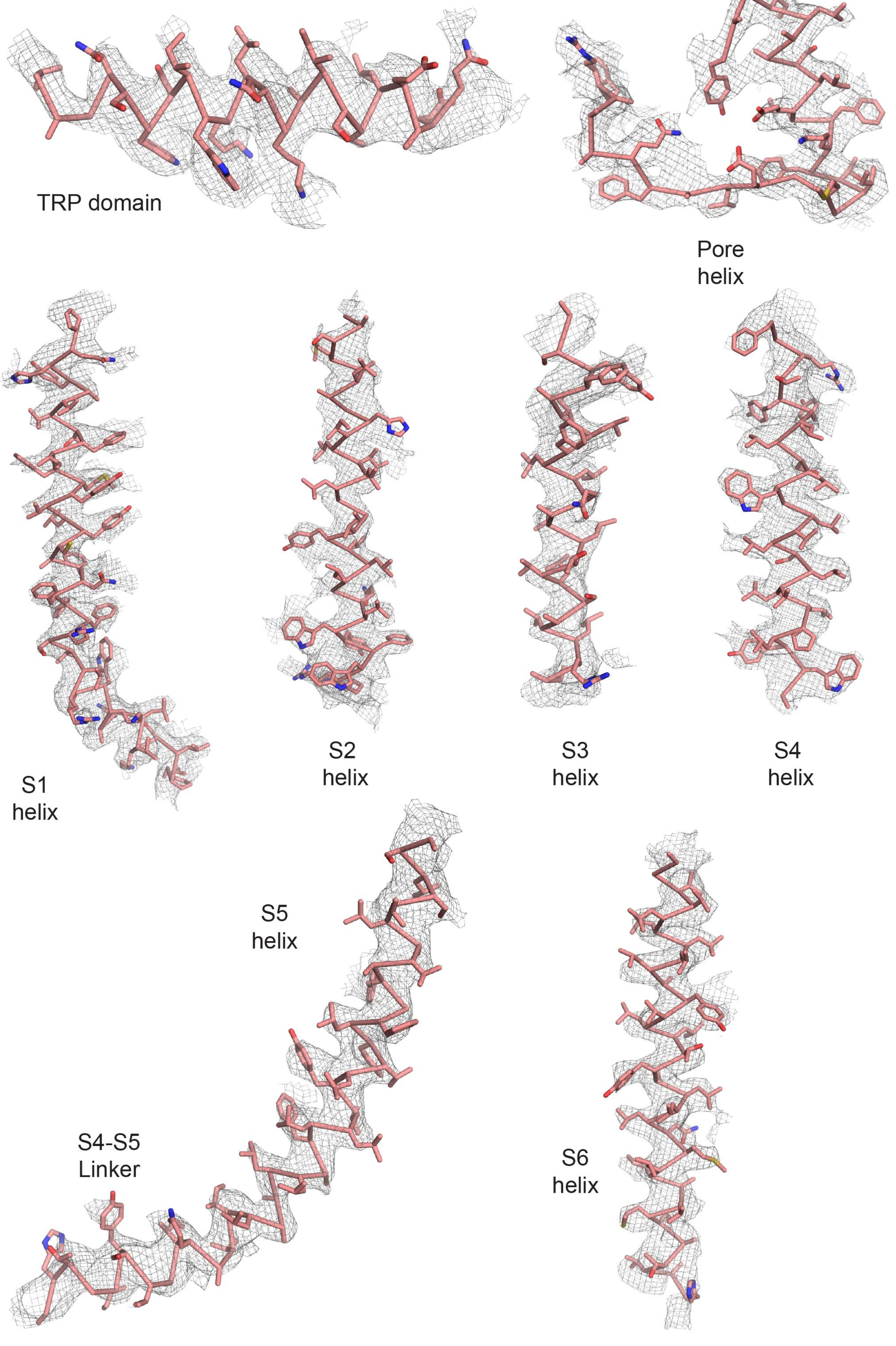

### Supplementary Figure 7

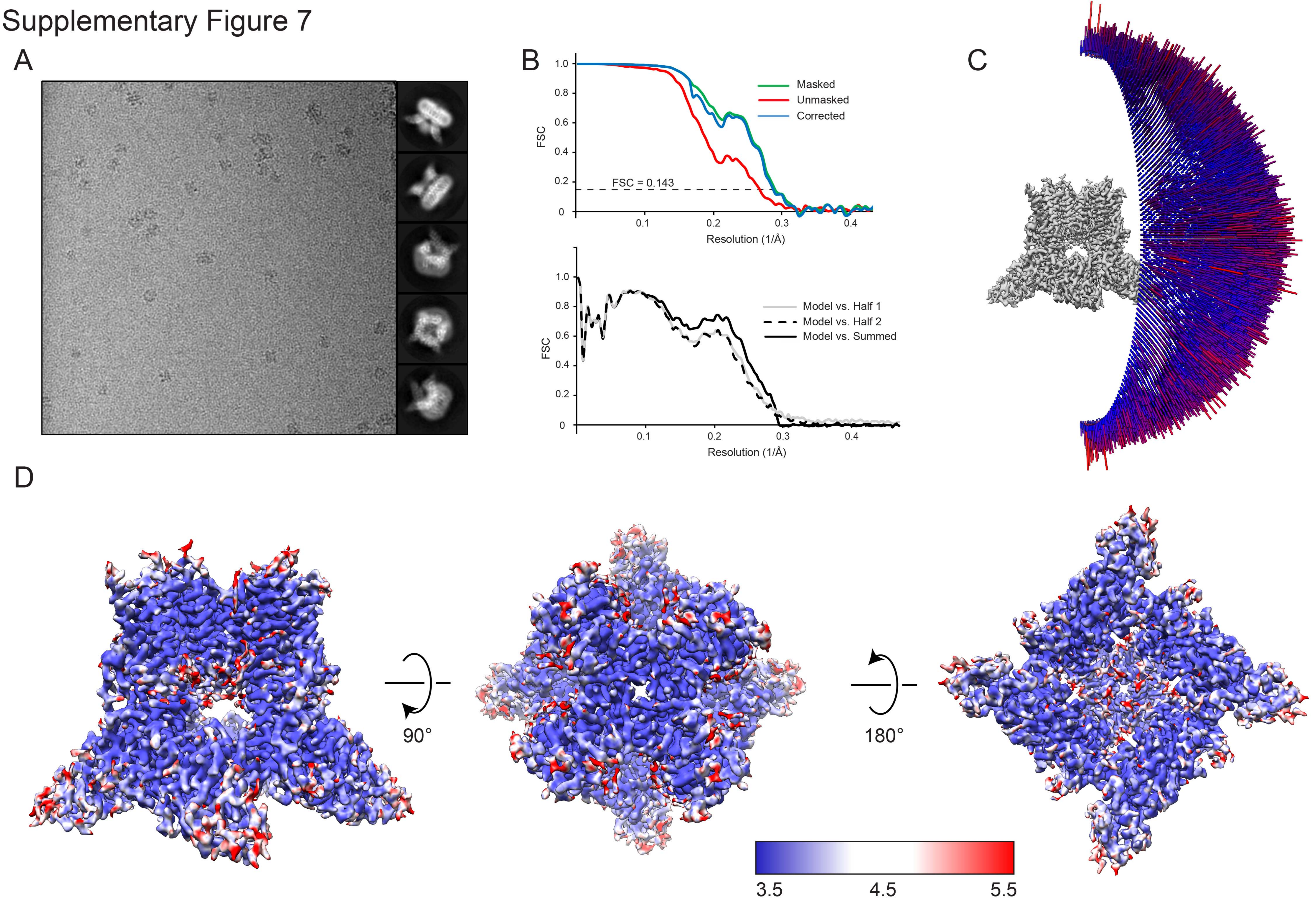

### Supplementary Figure 8

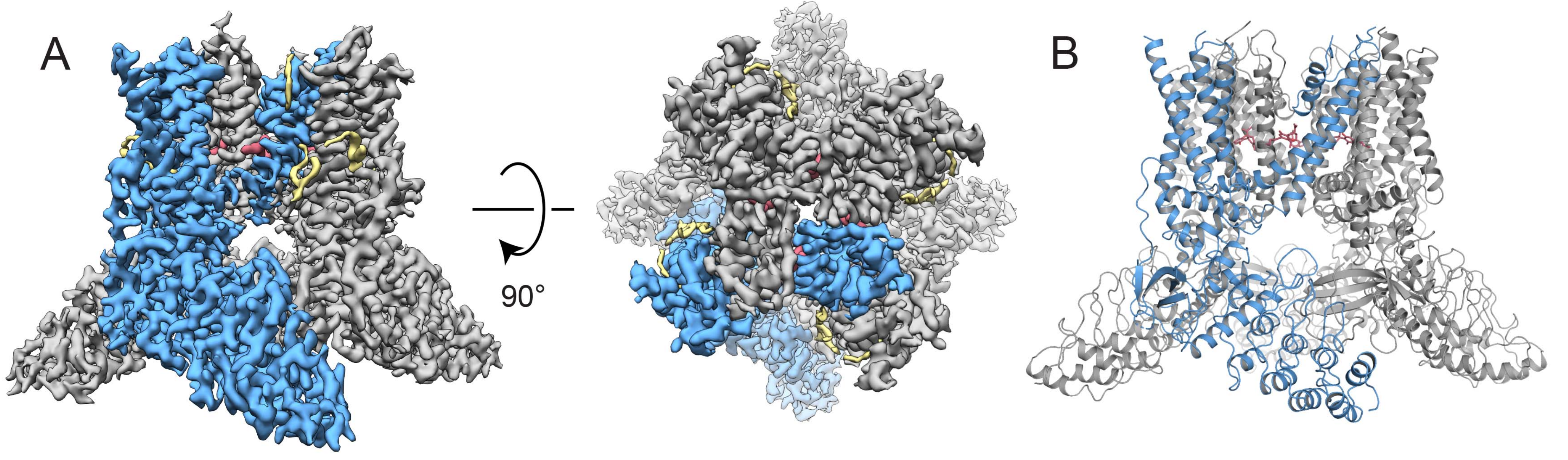

### Supplementary Figure 9

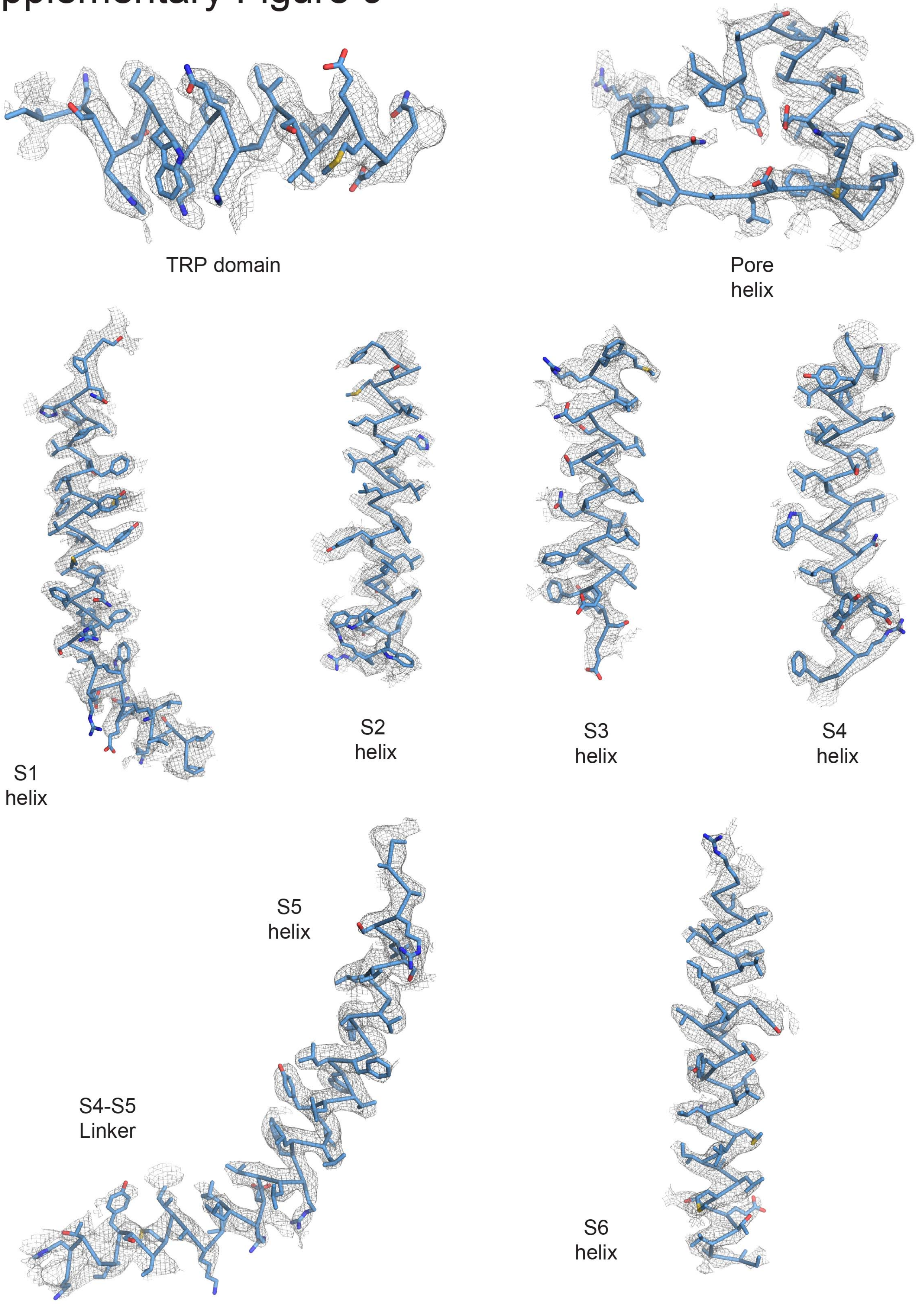

### Supplementary Figure 10

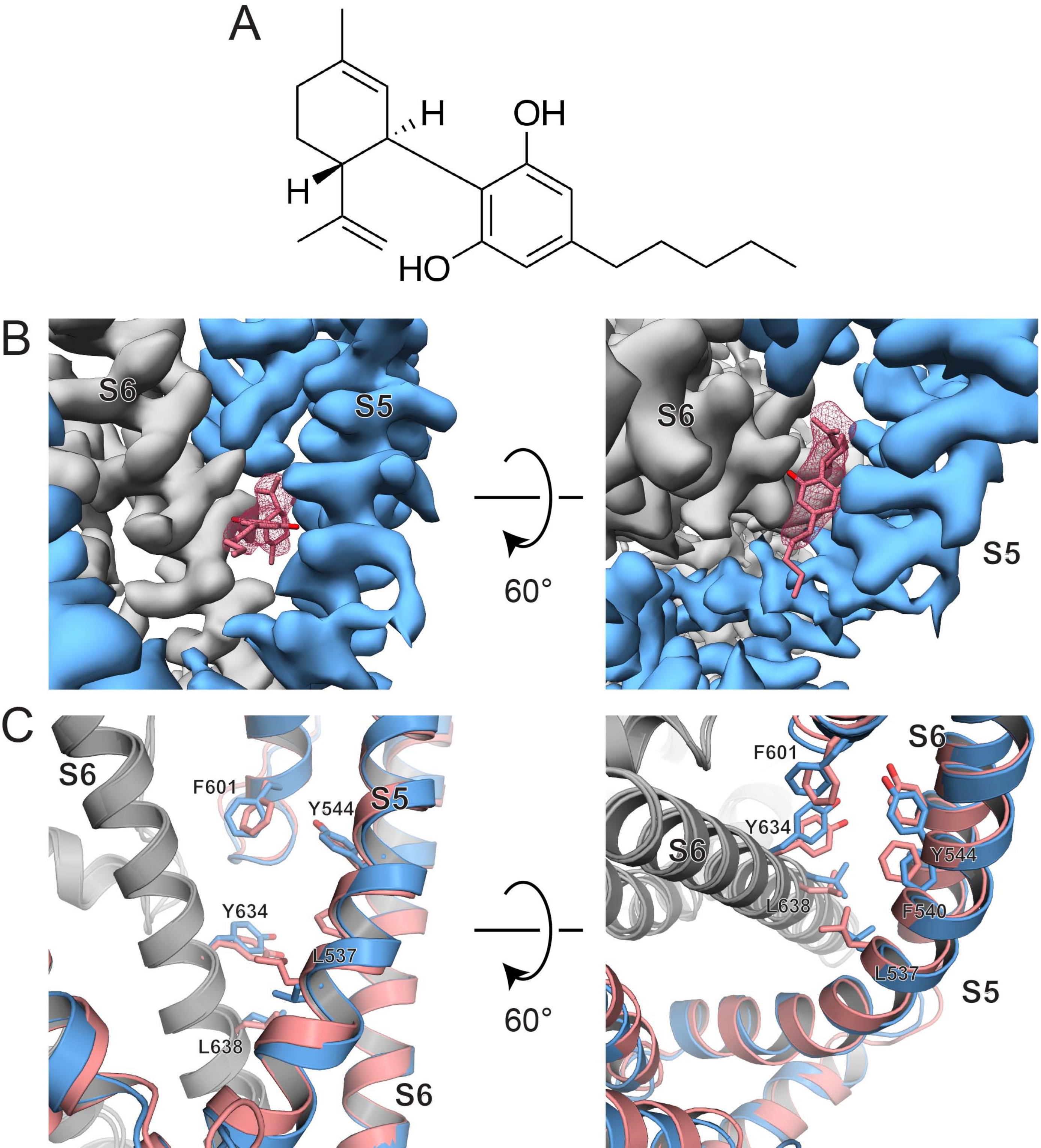

### Supplementary Figure 11

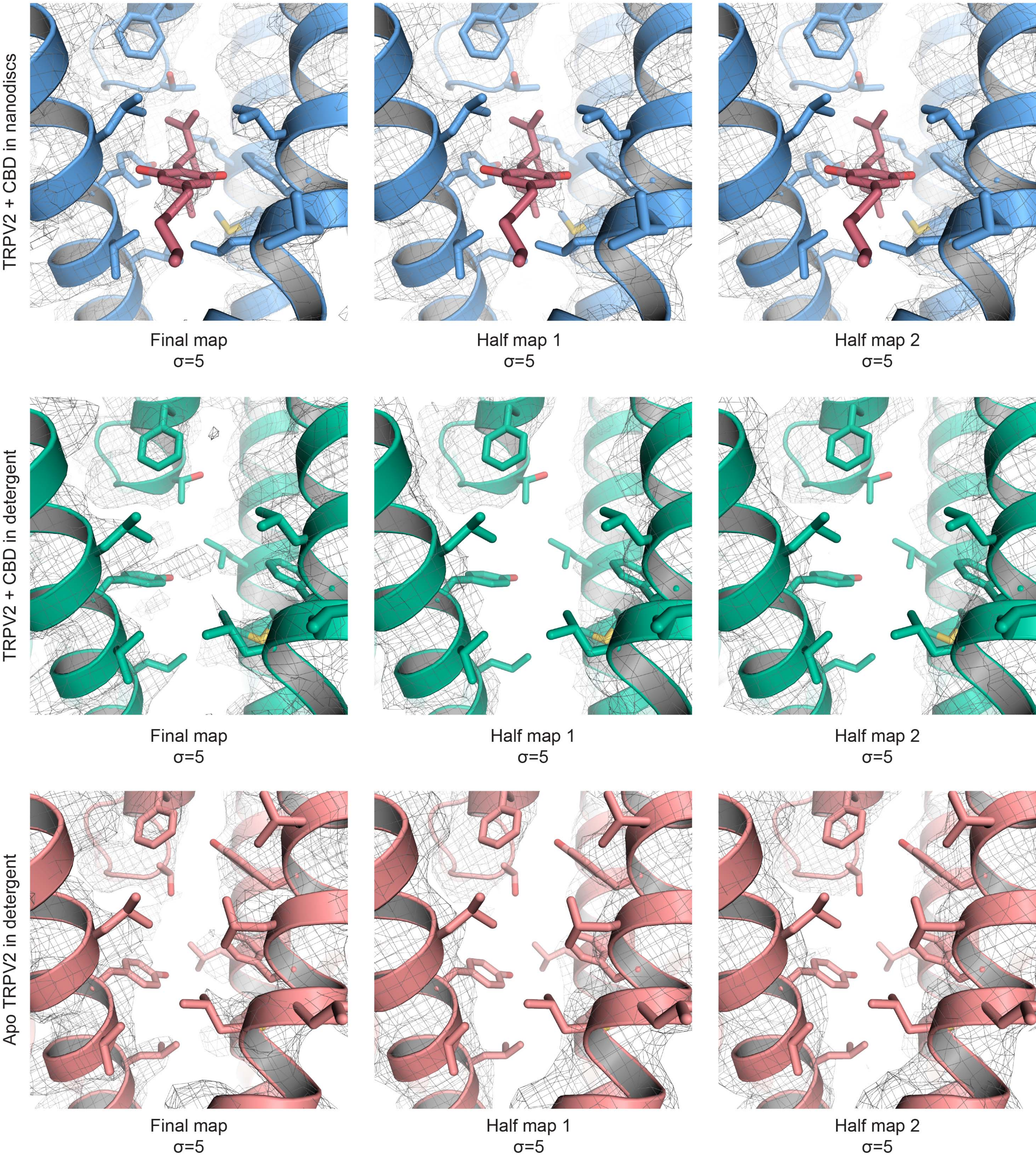

### Supplementary Figure 12

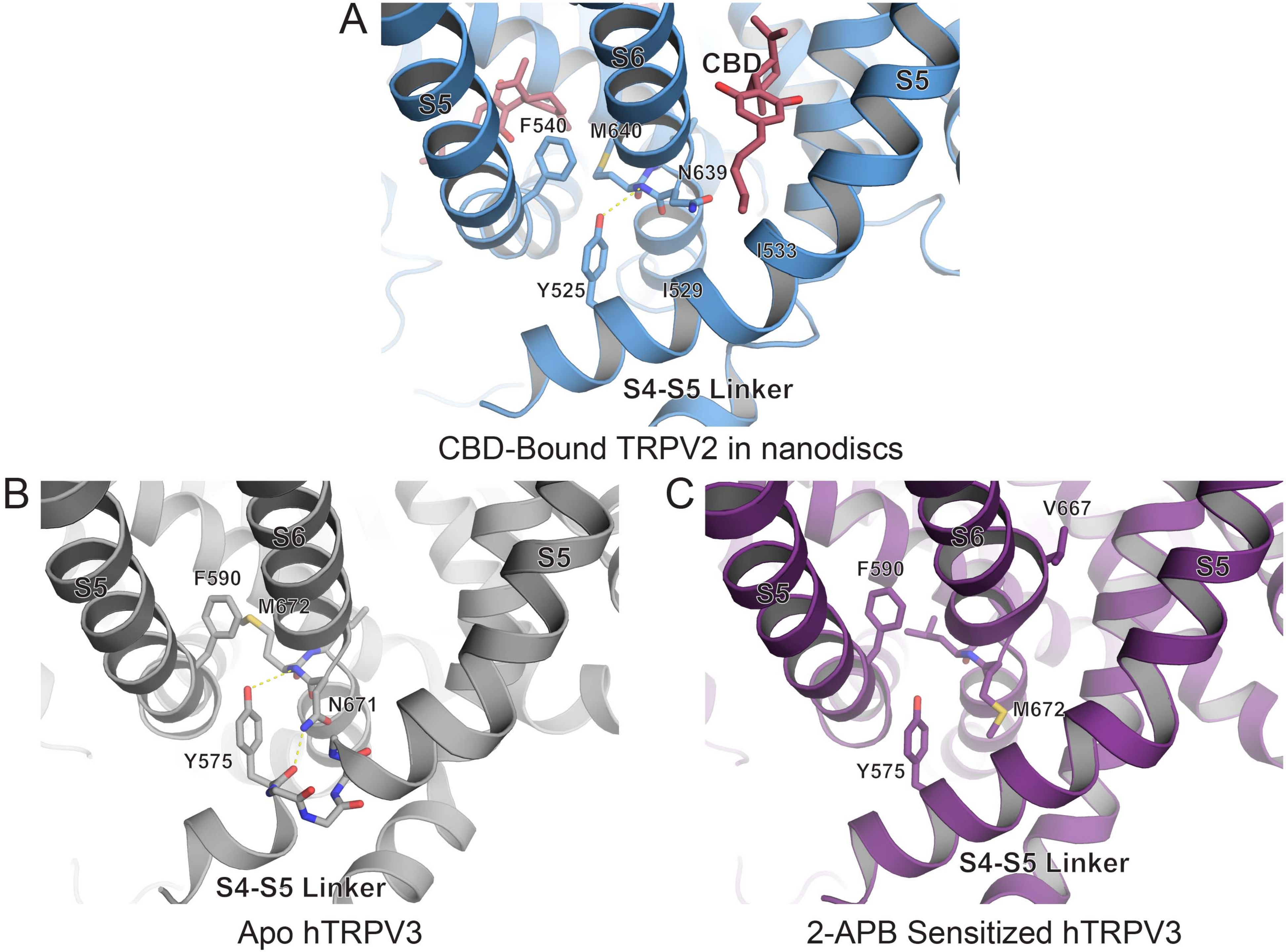

### Supplementary Figure 13

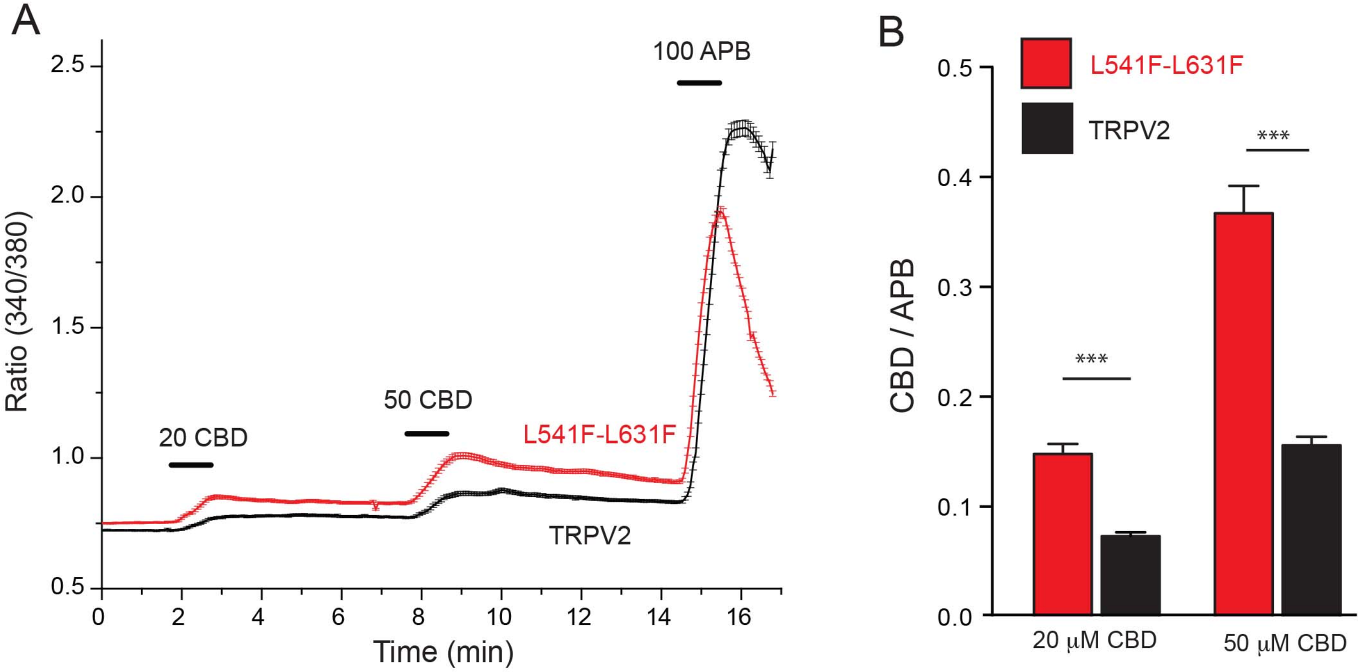

### Supplementary Figure 14

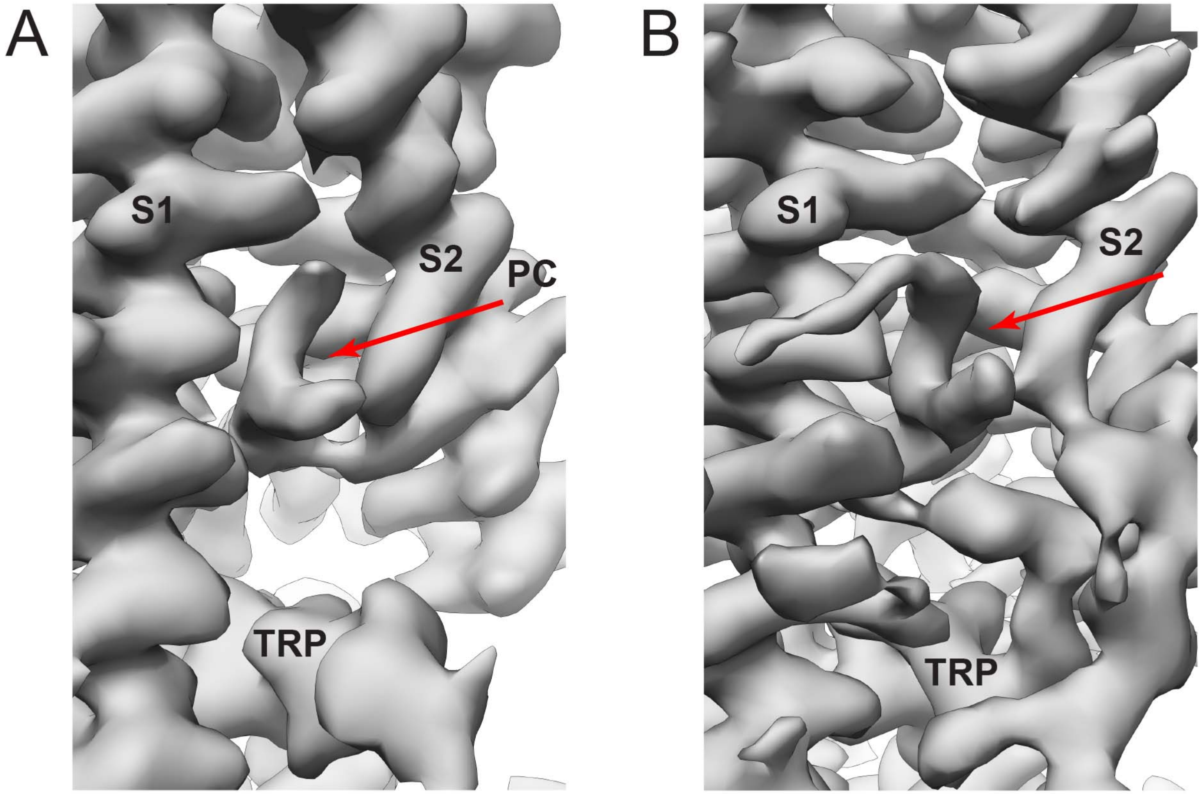

TRPV1 Apo in Nanodiscs (5IRZ)

TRPV2 + CBD in Nanodiscs

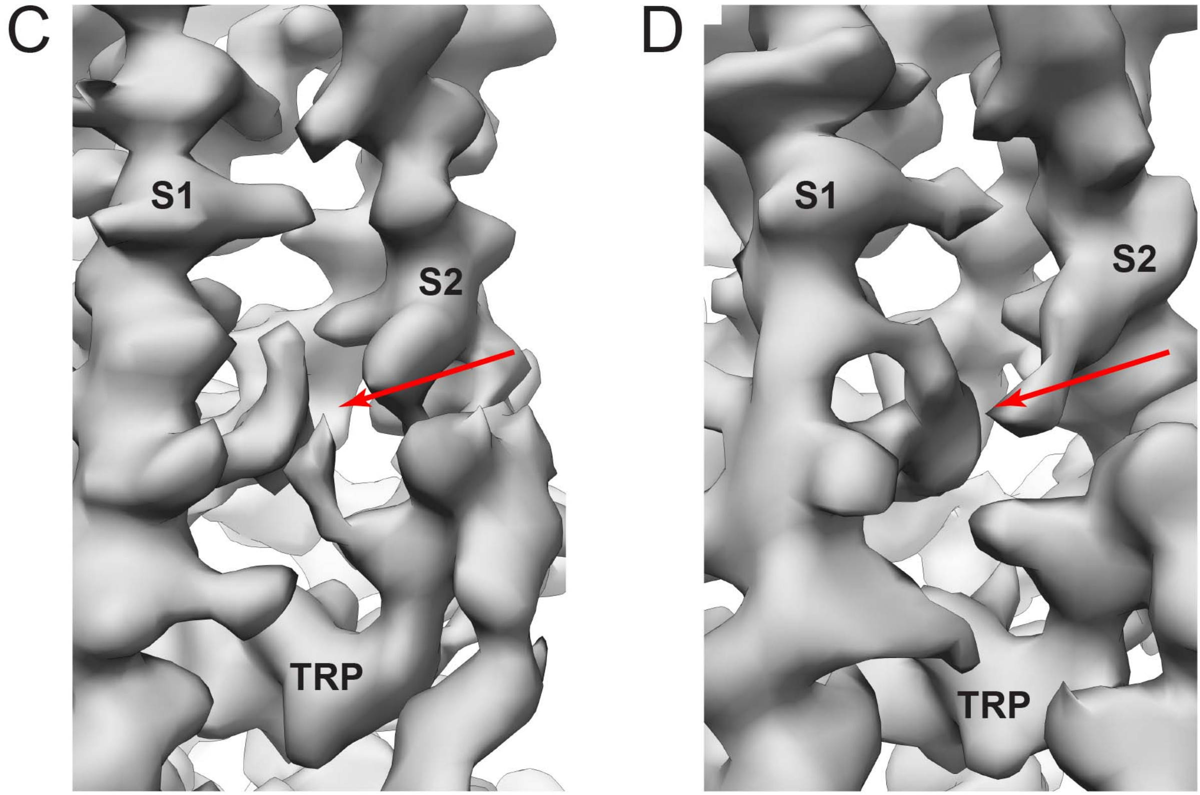

TRPV2 Apo in Detergent

TRPV2 + CBD in Detergent

#### Supplementary Figure 15

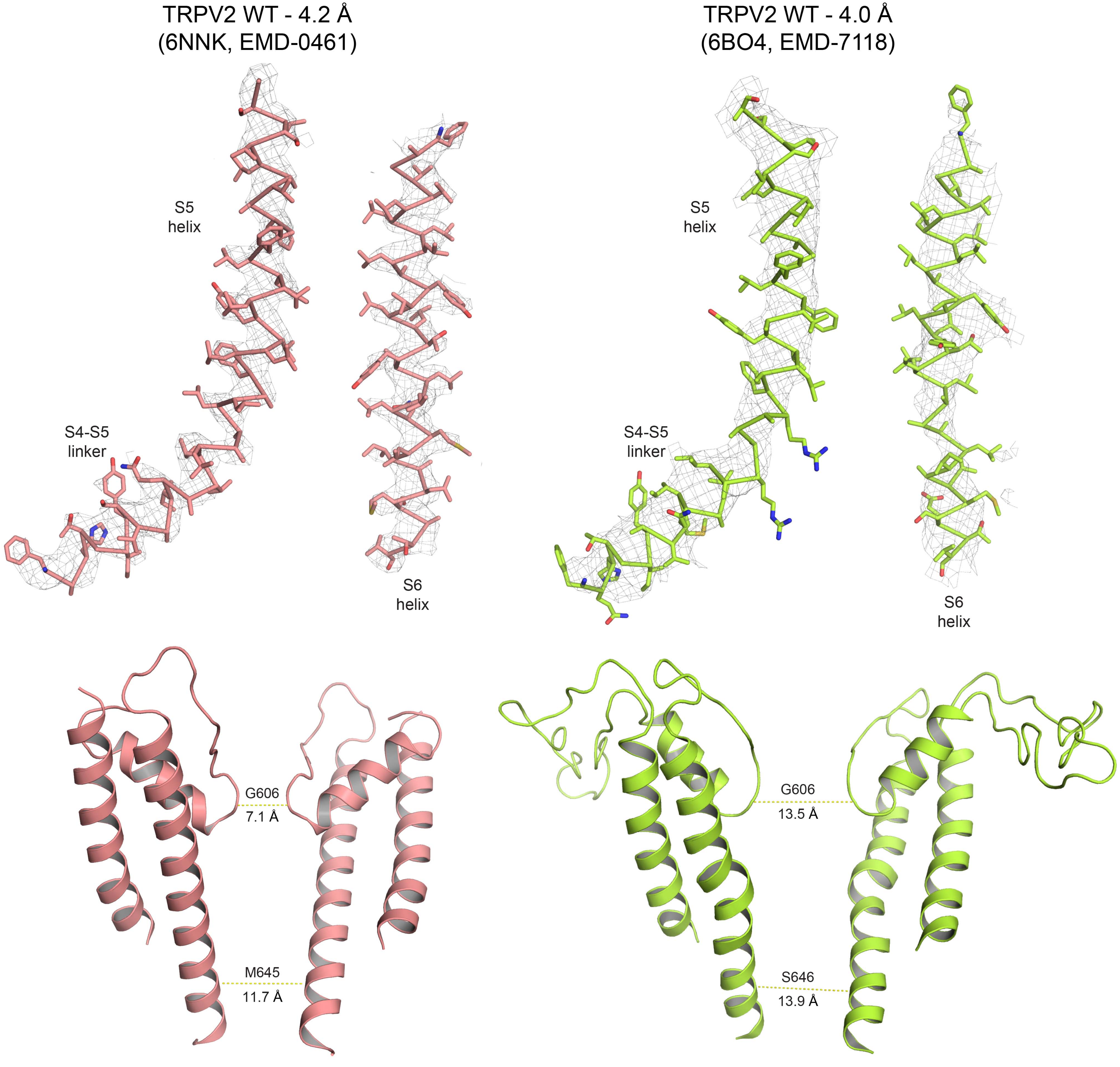

### Supplementary Figure 16

```
Residues marked: 534 537 538 540 541 544
Conservation
hTRPV1  KEYVASMVFSLALGWTNMLYYTRGFQQMGIYAVMIEKMILRDLCRFMFVYIVFLFGFSTAVVTLIEDGKNDSLPSES----TSHRWRGPACRPPDSSYNS
rTRPV1  KEYVASMVFSLAMGWTNMLYYTRGFQQMGIYAVMIEKMILRDLCRFMFVYLVFLFGFSTAVVTLIEDGKNNSLPMES----TPHKCRGSACKP-GNSYNS
hTRPV2  EWYLPLLVSALVLGWLNLLYYTRGFQHTGIYSVMIQKVILRDLLRFLLIYLVFLFGFAVALVSLSQEAWRPEAPTGPNATESVQPMEGQEDEGNGAQYRG
rTRPV2  EWYLPLLVLSLVLGWLNLLYYTRGFQHTGIYSVMIQKVILRDLLRFLLVYLVFLFGFAVALVSLSREARSPKAPEDNNSTVTEQPTVGQEEEP--APYRS
hTRPV3  KEYLACLVLAMALGWANMLYYTRGFQSMGMYSVMIQKVILHDVLKFLFVYIVFLLGFGVALASLIEKCPKDNKDCSS--------------------YGS
hTRPV4  EAYLAVMVFALVLGWMNALYFTRGLKLTGTYSIMIQKILFKDLFRFLLVYLLFMIGYASALVSLLNPCANMKVCNED------QTNCTVPTYPSCRDSET
hTRPV5  NGEVVPMSFALVLGWCSVMYFTRGFQMLGPFTIMIQKMIFGDLMRFCWLMAVVILGFASAFYIIFQTEDPTS--------------------LGQFYD
hTRPV6  SGEVVPMSFALVLGWCNVMYFARGFQMLGPFTIMIQKMIFGDLMRFCWLMAVVILGFASAFYIIFQTEDPEE--------------------LGHFYD

Residues marked: 601 604 631 634 635 637 638 640
Conservation
hTRPV1  LYSTCLELFKFTIGMGDLEFTENYDFKAVFIILLLAYVILTYILLLNMLIALMGETVNKIAQESKNIWKLQRAITILDTEKSFLKCMRKAFRSGKLLQVG
rTRPV1  LYSTCLELFKFTIGMGDLEFTENYDFKAVFIILLLAYVILTYILLLNMLIALMGETVNKIAQESKNIWKLQRAITILDTEKSFLKCMRKAFRSGKLLQVG
hTRPV2  ILEASLELFKFTIGMGELAFQEQLHFRGMVLLLLLAYVLLTYILLLNMLIALMSETVNSVATDSWSIWKLQKAISVLEMENGYWWCR-KKQRAGVMLTVG
rTRPV2  ILDASLELFKFTIGMGELAFQEQLRFRGVVLLLLLAYVLLTYVLLLNMLIALMSETVNHVADNSWSIWKLQKAISVLEMENGYWWCRRKKHREGRLLKVG
hTRPV3  FSDAVLELFKLTIGLGDLNIQQNSKYPILFLFLLITYVILTFVLLLNMLIALMGETVENVSKESERIWRLQRARTILEFEKMLPEWLRSRFRMGELCKVA
hTRPV4  FSTFLLDLFKLTIGMGDLEMLSSTKYPVVFIILLVTYIILTFVLLLNMLIALMGETVGQVSKESKHIWKLQWATTILDIERSFPVFLRKAFRSGEMVTVG
hTRPV5  YPMALFTTFELFLTVIDAPANYDVDLPFMFSIVNFAFAIIATLLMLNLFIAMMGDTHWRVAQERDELWRAQVVATTVMLERKLPRCLWPRSGICGCEFGL
hTRPV6  YPMALFSTFELFLTIIDGPANYNVDLPFMYSITYAAFAIIATLLMLNLLIAMMGDTHWRVAHERDELWRAQIVATTVMLERKLPRCLWPRSGICGREYGL
```

**Supplementary Table 1.** Cryo-EM data collection and model statistics

| | CBD-Bound TRPV2 in detergent (EMB-0462, PDB 6NNL) | CBD-Bound TRPV2 in Nanodiscs (EMB-0463, PDB 6NNM) | Apo TRPV2 in Detergent (EMB-0461, PDB 6NNK) |
|---|---|---|---|
| **Data collection and processing** | | | |
| Magnification | ~18,000 | ~45,500 | ~31,000 |
| Voltage (kV) | 300 | 300 | 300 |
| Defocus range (μm) | 1.4-3.0 | 0.8-3.0 | 1.5-3.0 |
| Pixel size (Å) | 1.38 | 1.07 | 1.29 |
| Symmetry imposed | C4 | C4 | C4 |
| Initial particle images (no.) | 820,953 | 685,702 | 420,081 |
| Final particle images (no.) | 49,496 | 25,206 | 51,045 |
| Map resolution (Å) | 4.3 | 3.5 | 4.2 |
| FSC threshold | 0.143 | 0.143 | 0.143 |
| Map resolution range (Å) | 3.5-5.5 | 3.5-5.5 | 3.5-5.5 |
| **Refinement** | | | |
| Model resolution cut-off (Å) | 4.3 | 3.5 | 4.2 |
| FSC threshold | 0.143 | 0.143 | 0.143 |
| Map sharpening *B* factor ($Å^2$) | -200 | -78 | -169 |
| Model composition | | | |
| Nonhydrogen atoms | 0 | 0 | 0 |
| Protein residues | 2488 | 2472 | 2420 |
| Ligands | 0 | 4 | 0 |
| R.m.s. deviations | | | |
| Bond lengths (Å) | 0.006 | 0.008 | 0.005 |
| Bond angles (°) | 0.938 | 0.692 | 0.635 |
| Validation | | | |
| MolProbity score | 1.70 | 1.83 | 2.17 |
| Clashscore | 4.00 | 6.49 | 11.69 |
| Poor rotamers (%) | 0.50 | 0.39 | 1.89 |
| Ramachandran plot | | | |
| Favored (%) | 91.30 | 92.45 | 94.51 |
| Allowed (%) | 8.70 | 7.55 | 5.49 |
| Disallowed (%) | 0.00 | 0.00 | 0.00 |
